# Natural variation in gene expression and Zika virus susceptibility revealed by villages of neural progenitor cells

**DOI:** 10.1101/2021.11.08.467815

**Authors:** Michael F. Wells, James Nemesh, Sulagna Ghosh, Jana M. Mitchell, Curtis J. Mello, Daniel Meyer, Kavya Raghunathan, Matthew Tegtmeyer, Derek Hawes, Anna Neumann, Kathleen A. Worringer, Joseph J. Raymond, Sravya Kommineni, Karrie Chan, Daniel Ho, Brant K. Peterson, Federica Piccioni, Ralda Nehme, Kevin Eggan, Steven A. McCarroll

## Abstract

Variation in the human genome contributes to abundant diversity in human traits and vulnerabilities, but the underlying molecular and cellular mechanisms are not yet known, and will need scalable approaches to accelerate their recognition. Here, we advanced and applied an experimental platform that analyzes genetic, molecular, and phenotypic heterogeneity across cells from very many human donors cultured in a single, shared *in vitro* environment, with algorithms (Dropulation and Census-seq) for assigning phenotypes to individual donors. We used natural genetic variation and synthetic (CRISPR-Cas9) genetic perturbations to analyze the vulnerability of neural progenitor cells to infection with Zika virus. These analyses identified a common variant in the antiviral *IFITM3* gene that regulated *IFITM3* expression and explained most inter-individual variation in NPCs’ susceptibility to Zika virus infectivity. These and other approaches could provide scalable ways to recognize the impact of genes and genetic variation on cellular phenotypes.

**HIGHLIGHTS:**

- Measuring cellular phenotypes in iPSCs and hPSC-derived NPCs from many donors
- Effects of donor sex, cell source, genetic and other variables on hPSC RNA expression
- Natural genetic variation and synthetic perturbation screens both identify *IFITM3* in NPC susceptibility to Zika virus
- A common genetic variant in *IFITM3* explains most inter-individual variation in NPC susceptibility to Zika virus

## INTRODUCTION

Humans and indeed all natural populations harbor immense diversity in biological traits, affecting almost all organ systems and physiological processes. Reservoirs of natural variation allow populations to adapt to existential crises and other strong selective pressures. Studies of quantitative traits and disease risk in populations – whether of humans, animals, or plants – reveal a large role for standing variation in shaping phenotypic diversity and enabling adaptation. And yet, the mechanisms by which natural genetic variation acts through molecular, cellular and developmental biology to shape trait variation are largely unknown.

Genetic variation in developmental processes has an important but still poorly understood role in shaping phenotypes throughout life. The spatial and cell-proliferative events that establish the cellular compositions of tissues – the cell types, their relative abundances, and their spatial arrangements – almost certainly affect those organs’ physiological properties years later. Consistent with this idea, many genetic discoveries in autism and schizophrenia point to genes that are most highly expressed during fetal brain development (Griesi-Oliveira et al., 2021; Mao et al., 2009; Stein et al., 2014). However, further understanding of the genetic and cell-biological mechanisms of such effects has been limited by the lack of experimental systems and clinical specimens relevant to the transient developmental window in which neural progenitor cells (NPCs) shape the cellular makeup of the brain; NPCs are no longer present when individuals’ phenotypes are ascertained or when tissue is sampled for analysis.

One mechanism by which genetic variants can lead to phenotypic diversity is through the regulation of gene expression, which has unknown – but almost certainly important – effects on signaling pathways, cell migration, and cell-cell interactions. Abundant human genetic variation has been ascertained and catalogued by the International HapMap Project (Belmont et al., 2005) and 1000 Genomes Project (Auton et al., 2015); collaborations such as the Genotype-Tissue Expression (GTEx; Aguet et al., 2017) project have identified thousands of expression QTLs (eQTLs) – associations of single nucleotide polymorphisms (SNPs) to RNA expression of nearby genes – in every adult organ analyzed. The relationships of these and other haplotypes to gene expression during development is largely unknown.

Relationships between common genetic variants and gene expression can also be revealed using *in vitro* stem cell-derived models, which have become increasingly adopted with advancements in differentiation protocols and the demonstrated similarities of these cells to corresponding cell types in tissues. Fast, reliable induction techniques for generating NPCs, neurons, glia, and oligodendrocytes from human embryonic stem cells (hESCs) and induced pluripotent stem cells (iPSCs) now enable investigations into gene expression variation. However, maintaining cells from many human donors in separate culture environments results in technical variation that can obscure biologically-relevant effects, and requires substantial resources and effort.

Though thousands of eQTLs have been found in every tissue, in iPSCs, and in iPSC-derived cells of various types, we know little today about the ways in which these effects percolate through cells to influence their functional phenotypes. Which of these genetic variants actually change a cell’s phenotype in a meaningful way? Biological pathways are built to be robust to many kinds of perturbation and may contain the effects of many genetic variants, even those that regulate a gene’s expression. The precise expression dosage of most genes is of unknown biological significance. Thus, it is essential to understand the relationships among genetic variants, gene expression, and the physiological phenotypes of cells that shape disease processes.

Here we describe the coordinated application of three approaches for dissecting these aspects of a cell’s life and their relationships to one another. One is genetic multiplexing, in which tens of thousands of cells from scores of donors are analyzed simultaneously by single-cell RNA-seq, revealing effects of genetic variants, cell-donor and cell-source properties on RNA expression at population scale. Another is “Census-seq”, a rapid, inexpensive method for relating cellular phenotypes to natural genetic variation by sequencing the genomic DNA from such “cell villages” (Mitchell et al., 2020). A third is functional CRISPR-Cas9 screens, which explore thousands of artificial genetic perturbations simultaneously (Joung et al., 2017; Ophir et al., 2014).

By applying these experimental platforms in a coordinated way, we find high levels of inter-individual variation in susceptibility of NPCs to the neurotropic Zika virus (ZIKV), then uncover a common, large-effect SNP in the antiviral *IFITM3* gene that explains more than 50% of this physiological variation and appears to act by regulating expression levels of *IFITM3* in NPCs. Analyses of the high-volume data from these same experiments helped us understand genetic, biological and technical sources of variation on gene expression in human pluripotent stem cells (hPSCs) and NPCs. The population-scale nature of these experiments made it possible to describe and quantify the effects of donor sex, cell source, and cell reprogramming methods. We also identify hundreds of NPC eQTLs, some of which affect expression levels of critical neurodevelopmental genes.

We hope that these results enable better understanding of sources of variation in hPSC-derived systems while suggesting possibilities for integrated experimental approaches that use natural and synthetic genetic perturbations to understand disease-relevant functional variants and their effects on specific cell types.

## RESULTS

### Analysis of cellular variation in “Dropulations”

To understand how natural genetic variation and donor-specific biological effects shape cellular phenotypes, we sought to eliminate other sources of variation by culturing cells from very many donors in a common environment and analyzing them all together. Transcribed SNPs contain abundant information that can be used to identify the donor of a biological sample using the alleles that are present in RNA transcripts. The combination of hundreds of transcribed SNPs can act as a cell-intrinsic barcode that uniquely identifies the donor of an individual cell (Kang et al., 2018). We further developed genetic multiplexing analysis: (i) to address challenges inherent to scRNA-seq experiments, such as ambient RNA; (ii) to utilize unique molecular indicators (rather than reads) as the basis for demultiplexing; and (iii) to allow scalability up to hundreds of potential donors (Methods). We provide the resulting software (“Dropulation”, for droplet-based sequencing of populations) in an open-source format (https://github.com/broadinstitute/Drop-seq).

We performed a variety of *in silico* and experimental analyses to confirm the accuracy with which cells were correctly assigned to donors by this approach (Methods, Fig. 1a, Supplementary Note, Fig. S1).

**Figure 1:**
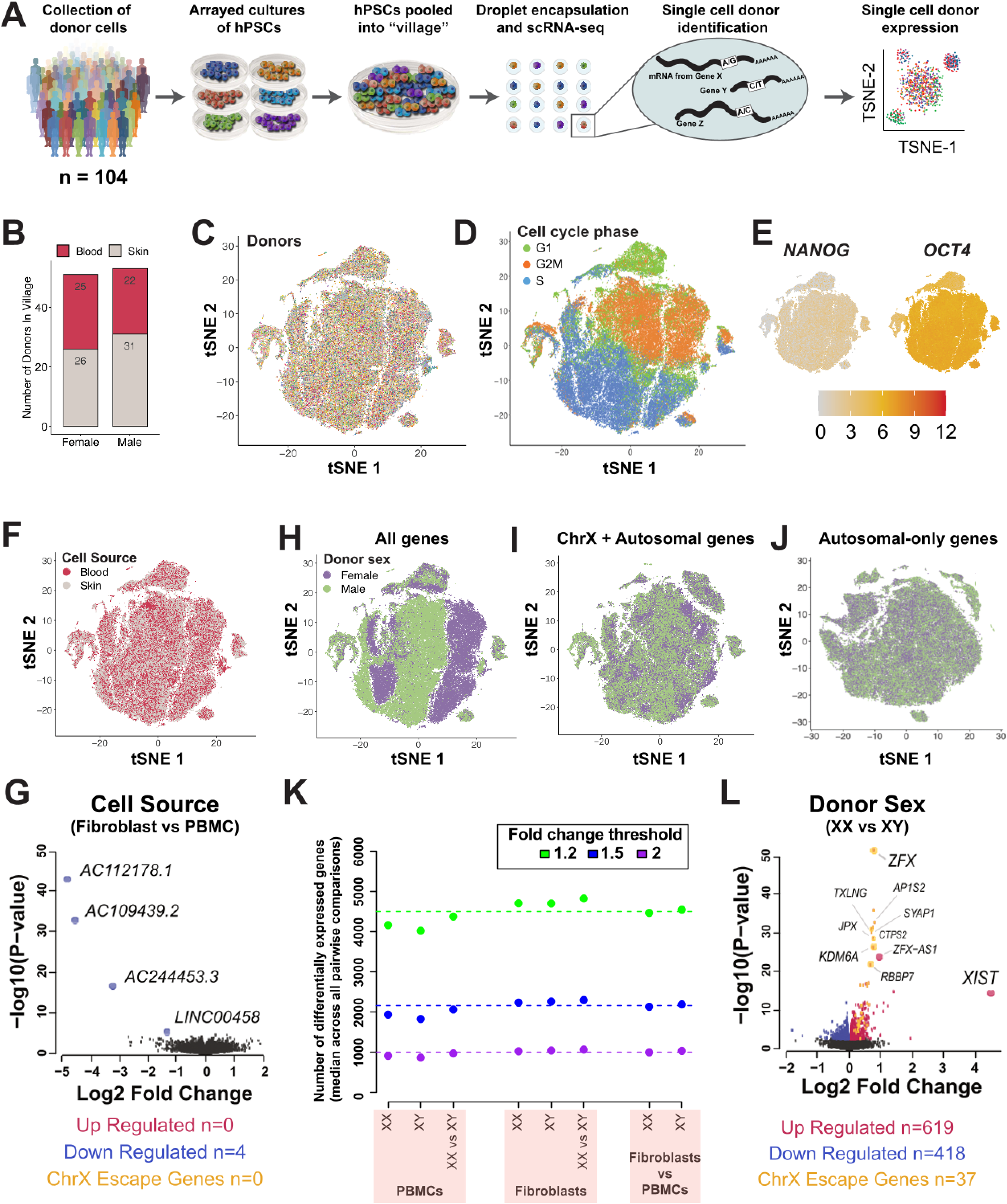
scRNA-seq characterization of human iPSC village. **A**, Schematic depicting cell village workflow. Human induced pluripotent stem cell lines from 104 donors were pooled into a single *in vitro* environment prior to scRNA-seq analysis; the donors of individual cells were identified from combinations of many (typically 200+) transcribed common SNPs. **B**, Composition of hiPSC village by donor sex and the tissue source of the cells used for reprogramming. **C-E**, Factors influencing variation in gene expression. (C) tSNE projection of scRNA-seq data color-coded by (C) donor of origin, as inferred from alleic information on scRNA-seq reads; and (D) cell cycle stage, as inferred from RNA expression levels. (E) Expression of pluripotency markers *OCT4* and *NANOG*. **F-G**, Modest effect of cell source (fibroblast vs. PBMC) is seen in (F) tSNE projection of the scRNA-seq data and (G) volcano plot of differential gene expression analysis grouped by donor cell source. **H-L**, Analyses of relationship of donor sex-chromosome status to RNA expression through a series of tSNE projections involving subsets of the scRNA-seq data consisting of (H) all genes, (I) X-chromosome plus autosomal genes, and (J) autosomal genes only. (K) Numbers of differentially expressed genes in pairwise comparisons of all iPSC donors, grouped by donor sex and cell source. Three different fold-change thresholds denoted by green (1.2-fold), blue (1.5-fold), and purple (2.0-fold) dotted lines. (L) Volcano plot of differential gene expression analysis grouped by donor sex.

The ability to measure mRNA expression in a common culture environment made it possible to quantify effects that have long been of great concern (but uncertain practical impact) in stem cell research, including the effects of donor sex and cell source. To characterize such effects, we established a 104-donor human iPSC “cell village” (Figure 1a) that included iPSCs derived from multiple cell sources (skin and blood cells) and from male and female donors (Figure 1b, Figure S2a). We profiled this cell village by scRNA-seq (10X Genomics platform) five days after pooling the 104 iPSC lines. We analyzed 86,185 cells by scRNA-seq, sampling their RNAs to an average depth of 104,160 UMIs per cell and assigning each cell to its donor-of-origin using Dropulation (Figure S2b-c).

When cultured together, the iPSCs from the 104 donors exhibited highly similar RNA-expression patterns, with cells from the 104 donors distributing similarly across a two-dimensional projection of the single cells’ RNA-expression patterns (Figure 1c). The primary source of variation in the individual cells’ expression profiles appeared to be their progress through the cell cycle (Figure 1d). Heterogeneity in cell state or identity can also impact gene expression (Figure S2d). As expected, the great majority of the hiPSCs expressed pluripotency markers *NANOG* and *OCT4* at high levels (Figure 1e); an exception (cluster 10, 4.3% of all cells) involved cells with higher-than expected expression of neural progenitor markers *RIPPLY3* and *SIX6*, suggesting that they have undergone spontaneous neural differentiation (Figure S2e). Another set of cells (clusters 6 and 7, 3.1% of all cells) exhibited high expression of *UTF1* and *MTG1*, suggesting poor differentiation potential; notably, these cells came from only a handful of cell lines (Figure S2f). A subset of single-cell expression profiles exhibited lower UMI counts and higher percentages of nascent transcripts with intronic reads (Figure S2g-h); we interpret these profiles to have arisen from nuclei (likely generated by unintentional cell lysis during handling) rather than intact cells.

iPSC lines are routinely created from skin (fibroblasts) or from blood (PBMCs); fundamental methodological questions involve the comparability of iPSCs created from different tissue sources. While it is reasonable to hypothesize that tissue source for iPSC reprogramming influences gene expression, our data suggests that such effects were small: Only four RNAs (all noncoding) exhibited genome-wide-significant differences in expression between the 56 skin-derived and 45 PBMC-derived iPSC lines (Figure 1fg; Data File 1). A more-sensitive enrichment analysis of sub-significant differential expression measurements indicated a modest enrichment associated with the somatic cell of origin (Data File 1): genes preferentially expressed in blood-derived iPSCs were nominally enriched in a myeloma related pathway, while genes preferentially expressed in skin-derived iPSCs were nominally enriched in a gene set associated with melanoma relapse (Figure S2i). This finding suggests that there is modest retention of epigenetic memory inherited from the parental cell source of origin in the iPS cell lines, but that few protein-coding genes are strongly affected by this memory.

Many human phenotypes show sex-biased differences, and RNA-expression levels of many genes differ on average between males and females in various tissues (Oliva et al., 2020). The extent to which cell-autonomous biology – as opposed to, for example, circulating hormones – contributes to such differences is unknown. The experimental design made it possible to isolate and quantify cell-autonomous effects, since (i) environmental and non-cell-autonomous effects were controlled across cells growing in the same culture, and (ii) very many male and female donors were sampled. The RNA-expression profiles of the individual cells initially appeared to be strongly distinguished by donor sex (Figure 1h), though this difference largely disappeared when we limited analysis to autosomal genes (Figure 1i-j), suggesting that it arose largely from Y-linked genes and genes that escape X chromosome inactivation.

How strong is sex-biased expression in iPSCs relative to the routine effects of inter-individual variation? We compared pairs of donors within and between sex to generate pairwise distributions of differentially expressed genes. We found similar numbers of differentially expressed autosomal genes in same-sex comparisons (XX vs XX; XY vs XY) as we did in across-group (XX vs. XY) comparisons, indicating that on average iPSCs from XX and XY individuals were roughly as different from each other as same-sex individual pairs regardless of cell source. These results suggest that differences in gene expression generated by donor sex and source cell-type are small compared to the routine effects of inter-individual variation, and that differences in expression of sex-chromosome genes do not lead to broader effects on cells’ biology in this context.

To further explore these modest, quantitative effects that become apparent in a population-scale analysis, we computed differential gene expression across all 104 donors using the limma/voom software package (Ritchie et al., 2015). As a point of reference, we performed similar analysis on reprogramming method cell source (Figure 1l). Consistent with earlier observations from tissue-level analysis (Oliva et al., 2020), effects were small (median log2FC upregulated genes = 0.15, downregulated genes = -0.10). To assess potential biological significance of the small transcriptional differences driven by cell source and donor sex, we performed gene set enrichment analysis using CAMERA (Wu and Smyth, 2012) and curated gene sets (C2) from the Molecular Signatures Database (MSigDB; (Subramanian et al., 2005). The most significant gene sets associated with differences across male and female donor lines mapped to pathways related to X chromosome inactivation and imprinting (Figure S2j). This signal disappeared when the analysis was limited to autosomal genes, indicating that differences in gene expression due to donor sex was largely driven by genes on the sex chromosomes (Data File 1).

The small effects of donor sex and cell source, relative to the routine effects of inter-individual variation, suggest that population-scale iPSC experiments can successfully utilize blood- and fibroblast-derived iPSCs together, though will ideally use designs in which these factors do not confound the effects of other variables of interest (such as genotype or case/control status).

### Effects of common genetic variation on RNA expression in hiPSCs

To map expression QTLs, we tested for associations between donors’ genotypes and their gene expression (summed across their individual iPSCs) using a linear regression model that included independent variables for biological and technical covariates, utilizing the MatrixEQTL analytical pipeline (Shabalin, 2012) (Figure 2a). Some 6,247 genes exhibited significant association to one or more SNPs (“eGenes”, at a q-value < 0.05) across the 104 hiPSC lines (Figure 2b, Data File 2).

**Figure 2:**
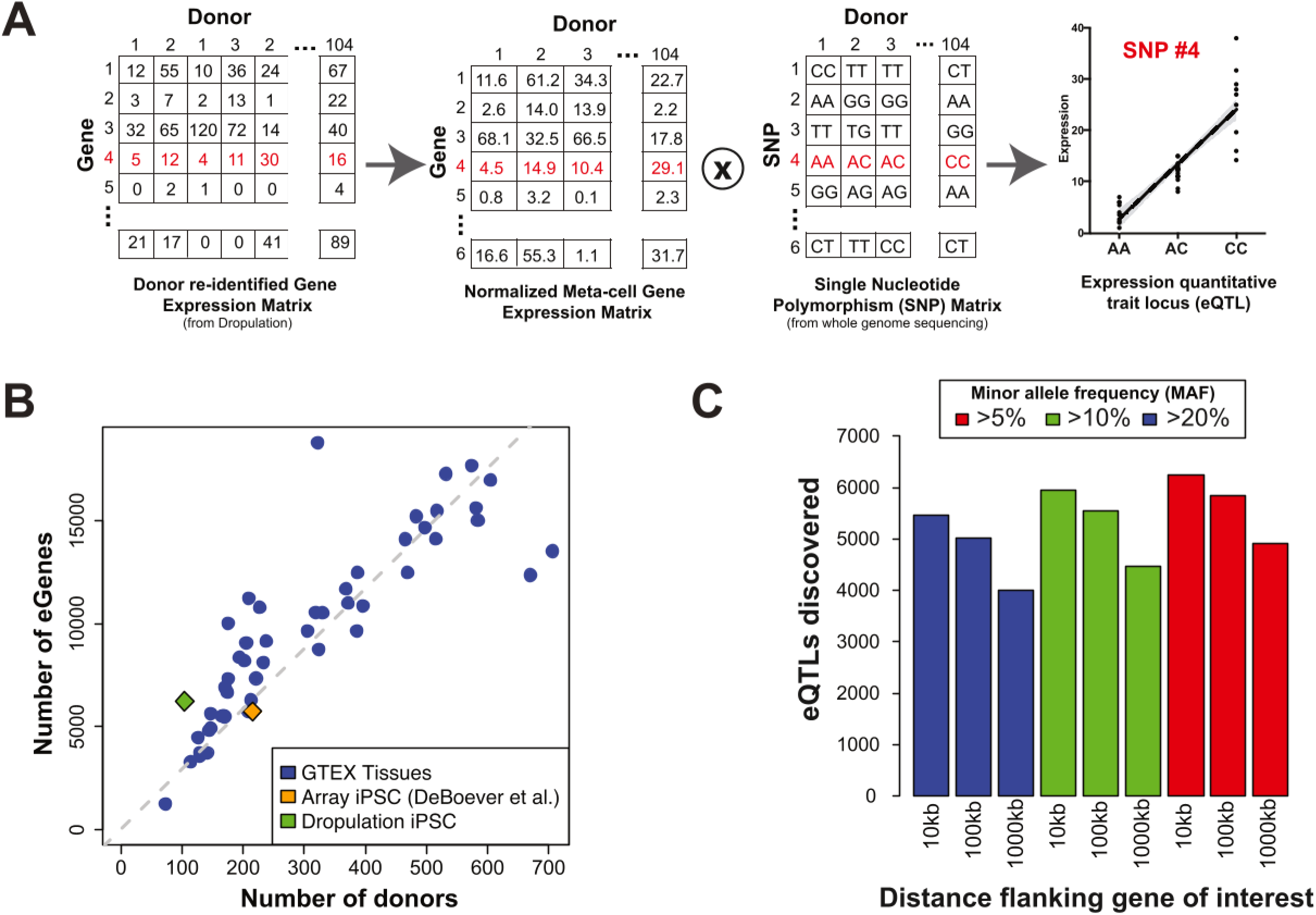
Detection of cis-expression QTLs (eQTLs). **A**, eQTL detection workflow. scRNA seq measurements for individual cells were summed into meta-cells (one per donor), which were then cross-referenced to SNP genotypes for eQTL analysis by MatrixEQTL. Red font denotes example data used for generating the eQTL plot for SNP#4. **B**, The number of eQTL-regulated gene (eGenes) detected by eQTL analysis in the 104-donor hiPSC village is shown in the context of other eQTL studies, including analyses (by GTEx) of 46 different human tissue types (blue circles) and earlier analysis of hPSCs in an arrayed format (orange squares). **C,** Optimization of eQTL search parameters including search windows (genomic distance searched around each gene) and minor allele frequencies (MAF).

To critically evaluate the utility of the genetically multiplexed format for eQTL discovery, we compared rates of eQTL detection across studies of diverse tissue and cell types, including GTEX and a report discovering eQTLs in human stem cells that were cultured and analyzed in an arrayed format (DeBoever et al., 2017). Across these studies, power to discover eQTLs increased almost linearly in relation to the number of donors sampled (with a slope of 29.2 eQTLs per donor). Analysis of cells in a “cell village” format identified 60 eQTLs per donor (Figure 2b), suggesting that the multiplexed approach helped increase the sensitivity of eQTL detection, likely by controlling for technical and environmental factors that introduce noise into most eQTL discovery efforts.

To assess the relationship of these iPSC eQTLs to eQTLs discovered by large-scale analysis of human tissues, we performed a sign test, asking what fraction of these SNPs exhibited the same direction of association to the same gene-expression phenotypes in other tissues and experiments. We found an 88% concordance (by sign test) between the eQTLs we found, and eQTLs found by the arrayed analysis of many individual iPSC lines. In addition, we observed similar sign test results when comparing to GTEx tissue and cell type data, with the strongest concordance coming from tissues with many mitotic cells (Figure S3a).

Finally, we explored the significance of eQTL search parameters, such as the search window (around each gene) used for cis-eQTL discovery, and the allele frequencies of the SNPs considered; both involve a tradeoff between the increased sensitivity of a broader search, and the reduction in sensitivity when multiple hypothesis testing is fully addressed in such a search (Figure 2c). We hope that this information is useful in guiding other eQTL discovery studies.

### Differential susceptibility to Zika virus infection across donor lines

hPSCs and hPSC-derived cell types offer a unique opportunity to analyze genetic influences upon the physiological phenotypes of living cells. They also present an opportunity to learn from both natural genetic variation and synthetic genetic perturbations. We sought to identify and understand genetic effects upon a physiological phenotype, the vulnerability of human NPCs to the neurotropic Zika virus.

Interactions with viruses have significantly altered the human genetic landscape in the form of an arms race between pathogen and host (Enard et al., 2016). Several recent reports have chronicled SNP associations with clinical outcomes of influenza, HIV, and SARS-CoV-2 viral infections, among others (Helminen et al., 1999; McLaren et al., 2015; Zhang et al., 2020, 2013). Many of these SNPs may affect the expression of human genes that play prominent roles in viral entry or replication mechanisms, as well as those that mediate the innate immune response (Kenney et al., 2017), and in doing so influence the probability of productive infection and spread to neighboring host cells. While numerous relationships between specific SNPs and viruses have been identified across a range of cell types, our understanding of the interplay among human genetic variation, viral susceptibility, and the brain is still in its infancy.

The Zika virus is a global health concern, with an outbreak that originated in South America and spread to over 50 countries between 2015 and 2016 (Barbeito-Andrés et al., 2018; Rossi et al., 2018; Saxena et al., 2018). While most healthy adults are asymptomatic after infection, prenatal exposure to this mosquito-borne virus can result in Congenital Zika Syndrome (CZS), which is characterized by severe neurological outcomes including microcephaly and neurodevelopmental delay (Brasil et al., 2016; Cauchemez et al., 2016; Mlakar et al., 2016). Interestingly, only 30% of prenatal infections during this outbreak resulted in CZS, and 95% of these cases were reported in only one country—Brazil (Nielsen-saines et al., 2019; de Oliveira et al., 2017). While many factors have been hypothesized to explain these observations, including maternal nutrition and socioeconomic status (Barbeito-Andrés et al., 2020; Souza et al., 2018), other potential factors – including human genetic diversity – can contribute to differential responses to this viral pathogen (Borda et al., 2021; Gomes et al., 2021; Santos et al., 2019).

ZIKV infection leads to CZS primarily through its preferential targeting of proliferating NPCs and glial cell types over post-mitotic neurons (Retallack et al., 2016; Tang et al., 2016). We therefore used the <u>S</u>tem cell-derived <u>N</u>GN2-<u>a</u>ccelerated Progenitor (SNaP) *in vitro* model of human dorsal telencephalic neural progenitors, which we recently showed are capable of supporting the complete ZIKV life cycle (Wells et al., 2018), to investigate ZIKV pathogenesis in a microcephaly-relevant cell type. To quantify inherent differences in ZIKV susceptibility *in vitro*, we infected 24 independently-cultured hESC-derived SNaP lines with ZIKV-Ug (MOI = 1) and measured infectivity levels at 54 hpi (Figure 3a). We observed a surprisingly high degree of variability in viral sensitivity, with mean infectivity rates ranging from 0.7% (RUES1 cell line) to 99.4% (Mel2; Figure 3b).

**Figure 3:**
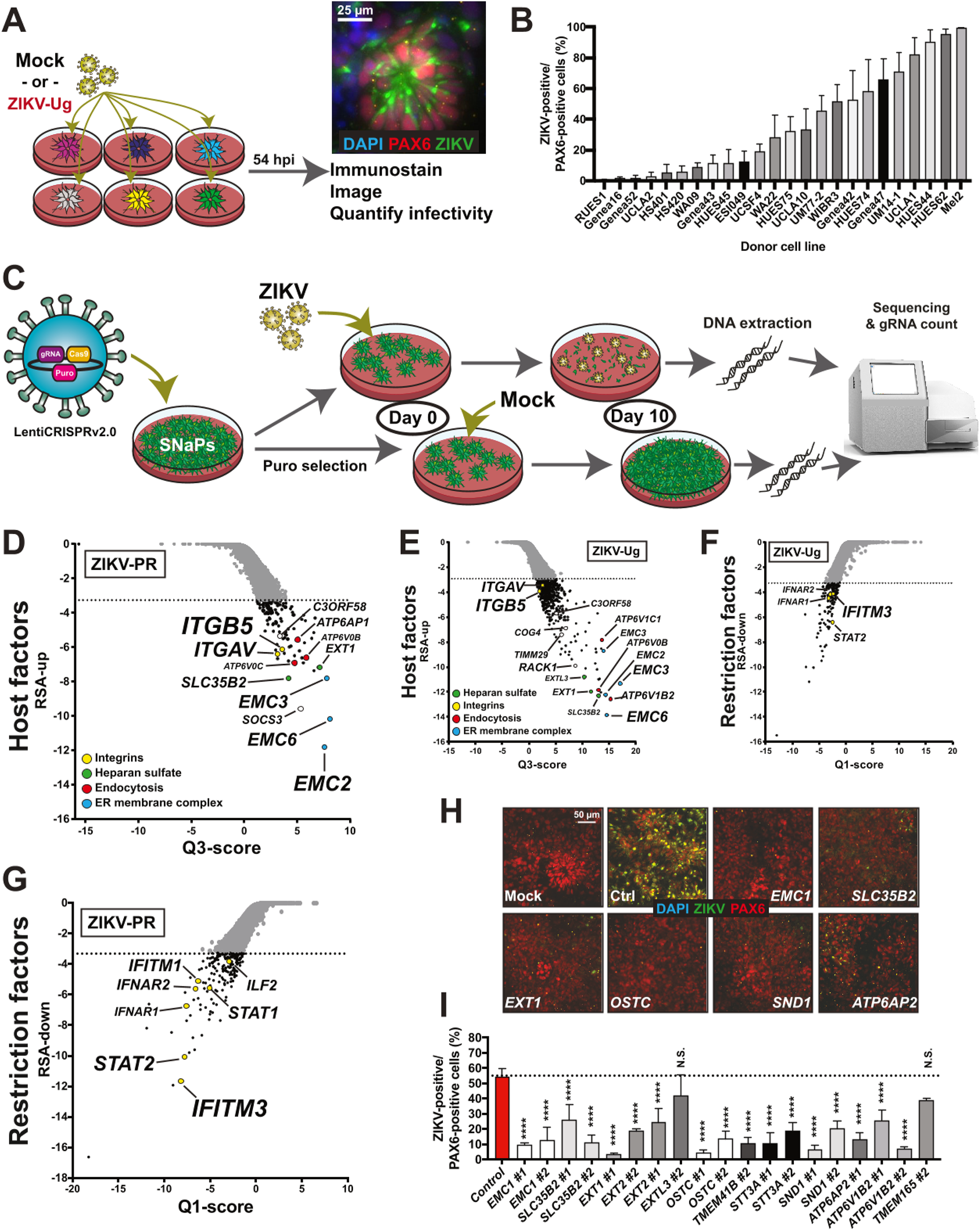
ZIKV infectivity varies across human NPC donor lines. **A**, Schematic depicting arrayed infection of 24 hESC-derived SNaP lines with ZIKV-Ug (MOI = 1). Representative immunostaining image of ZIKV-Ug 4G2 envelope protein at 54 hpi. Scale bar = 25 µm. **B**, Quantification of arrayed infectivity assays (ZIKV-Ug, MOI = 1) at 54 hpi. **C**, Design of whole-genome CRISPR-Cas9 screens. SW7388-1 hiPSCs were induced to SNaPs before transduction of the Brunello gRNA library via the LentiCRISPRv2.0 system. SNaPs used for the ZIKV survival screen were infected with ZIKV-Ug (MOI = 1), ZIKV-PR (MOI = 5), or mock infection media, and harvested 10 days later for DNA sequencing. ZIKV-infected samples were compared to Day 10 mock controls to identify genes that enhance ZIKV survival. **D-E**, RSA plots depicting gene level results of the enriched host factor genes from (D) ZIKV-PR and (E) ZIKV-Ug screens. Dashed line denotes hit cutoff for genes enriched in Day 10 ZIKV samples relative to mock controls (adjusted p-value < 0.05). **F-G**, RSA plots depicting gene level results of the depleted restriction factor genes from (F) ZIKV-PR and (G) ZIKV-Ug screens. Dashed line denotes hit cutoff for genes depleted in Day 10 ZIKV samples relative to mock controls (adjusted p-value < 0.05). **H-I**, Confirmation of primary screen hits. H1-Cas9 SNaPs were transduced with individual gRNAs, expanded, and exposed to ZIKV-Ug (MOI = 1). At 54hpi, infectivity was measured relative to mock controls. (H) Representative images and (I) quantification. Dashed line denotes infectivity levels for non-targeting gRNA controls. One-way ANOVA with Dunnett’s test for multiple comparisons. Data presented as mean ± SD. N.S. = not significant, ****p<0.0001.

These data suggest that cell-intrinsic factors make a large contribution to the inter-individual variation in vulnerability to ZIKV. We sought to understand this through both natural genetic variation and synthetic genetic perturbations, which we hypothesized might lead us to the same genes.

### Genome-wide CRISPR-Cas9 screens reveal ZIKV host factors in human NPCs

Viruses commandeer cellular proteins and pathways at each stage of their life cycle in an effort to propagate and spread to additional cells. The exact host biological processes subjected to these manipulations differ based on virus and cell type. Genome-wide screens have identified ZIKV host genetic factors in various cell types (Savidis et al., 2016a; Wang et al., 2020), including hPSC-derived NPCs (Li et al., 2019). We wanted to run similar genome-wide CRISPR-Cas9 positive selection survival screens in SNaPs to determine if the same genes are important for ZIKV infection of our cellular model, as well as to limit the number of genes that could potentially explain the differential ZIKV susceptibility we observed across donor lines.

We transduced SW7388-1 SNaPs with the Brunello human CRISPR knockout pooled library packaged in the all-in-one (Cas9 + gRNA) LentiCRISPRv2.0 vector (Figure 3c). LentiCRISPR-expressing SNaPs underwent 7 days of puromycin selection and expansion prior to infection with ZIKV-Ug (MOI = 1), ZIKV-PR (MOI = 5), or mock-infection media. Ten days later, all samples were harvested and prepared for DNA extraction and next generation sequencing (NGS) before the number of gRNA reads in each sample was counted and used for subsequent comparative analysis. Our results showed high gRNA coverage (<0.15% missing gRNAs across replicates), a high correlation among replicates (Pearson r values >0.92), and the expected limited effects of NT gRNA controls (Figure S4a-b; Data File 3).

The enrichment of sequencing reads for a gRNA targeting a particular gene in the ZIKV-infected samples relative to mock controls at Day 10 would suggest that disruption of said gene confers protection from ZIKV-mediated cell death. Using the redundant siRNA activity (RSA) statistical method (König et al., 2007), we detected 102 and 765 significantly-enriched candidate host factor genes in the ZIKV-PR and ZIKV-Ug samples, respectively, relative to mock controls (adjusted p-value < 0.05; Figure 3d-e). Despite the disparity in the numbers of significantly enriched genes, which is likely due to the higher noise associated with the reduced levels of cell death in the ZIKV-PR screen (ZIKV-Ug: 3.83% cell viability, ZIKV-PR: 26.0% cell viability; Data not shown), the two screens otherwise showed strong concordance: 75% (77/102) of the ZIKV-PR hits were also genome-wide hits in the ZIKV-Ug screen (Data File 3), indicating that the mechanisms that mediate ZIKV-induced NPC death are grossly similar between the two strains.

As expected, the strongest-effect genes and pathways from the survival screens corresponded to genes involved in known flaviviral infection and replication mechanisms, including heparan sulfate biosynthesis (*EXT1, SLC35B2, B4GALT7, EXT2*), endosomal acidification (*ATP6AP2, ATPV1B2, ATP6V0B*), the oligosaccharyltransferase (*OSTC* and *STT3A*) and endoplasmic reticulum membrane complexes (*EMC1*-*7*), and the oligomeric Golgi complex (*COG3, COG4, COG8, COG1*). Many of these genes were detected in the whole-genome experiments performed using conventional hPSC-derived NPCs (Li et al., 2019). Our screen identified additional genes not previously attributed to ZIKV pathogenesis, including the viral translation regulator *RACK1* (Majzoub et al., 2014) and the mitochondrial translocase *TIMM29*, thus highlighting inhibitors of these proteins as potential therapeutics. Previously nominated ZIKV entry factors *AXL, TYRO3, MERTK, CD209, HAVCR1*, and *TIMD4* were not significantly enriched in our primary screen, consistent with our previous findings (Wells et al., 2016). Our screen did however highlight the *ITGAV* and *ITGB5* genes, which together encode the transmembrane αVβ5 protein and have previously been shown to serve as an entry factor for other viruses (Ballana et al., 2011).

We also detected 195 and 147 genes that were significantly depleted in the ZIKV-PR and ZIKV-Ug screens, respectively, suggesting that their ablation rendered cells more sensitive to ZIKV-mediated cell death (adjusted p-value < 0.05; Figure 3f-g). The 48 depleted genes that were detected in both screens included the Type I interferon-responsive genes *IFNAR1-2, STAT2*, and *IFITM3*.

The results of the primary survival screen were confirmed through CRISPR- and antibody-mediated secondary assays. We first generated a H1 hESC line that constitutively expresses Cas9 (clone H1-36-23) and validated its genome-editing properties (Figure S4c-f). H1-Cas9 hESCs were then induced to SNaPs and transduced with lenti-gRNAs in an arrayed format for infectivity assays at 54 hpi and cell viability assays at 120 hpi using ZIKV-Ug (MOI = 1). We targeted a subset of genes that were significantly enriched in both SNaP survival screens and represented the spectrum of protein complexes and biological processes identified by these primary screens. As predicted, administering guides that targeted the EMC genes, *SLC35B2*, OST complex components, and vATPase subunits, but not TAM receptors, resulted in significantly improved cell viability and reduced infectivity compared to NT gRNA controls (Figure 3h-i, S4g-i). Interestingly, disruption of the transmembrane-spanning integrin subunit ITGB5 improved cell viability and dramatically reduced levels of infection (Figure S4j). Similar protection was observed when SNaPs were treated with anti-integrin αVβ5 antibody prior to ZIKV-Ug exposure (Figure S4k-l) in agreement with two recent reports that also nominated integrin αVβ5 as a ZIKV entry factor in neural cells (Wang et al., 2020; Zhu et al., 2020).

These survival screens implicated the host genes appropriated by ZIKV along every step of its lifecycle, as well as the antiviral effectors utilized by NPCs to fight infection. The outcomes of these screens are largely consistent with previous findings in hPSC-derived NPCs generated using standard induction protocols (Li et al., 2019), providing additional confirmation that SNaPs behave like NPCs. Along with the discoveries from recent large-scale drug screens in human NPCs (Xu et al., 2016; Zhou et al., 2017), our results should be taken into consideration when developing novel clinical therapies against CZS, for which there are currently no approved treatments. Perhaps most importantly, these data nominate a small number of genes as potential mediators of the differential viral susceptibility we observed across SNaP lines.

### Identification of expression QTLs in ZIKV host factors

Having identified (in the CRISPR screens) genes that mediate ZIKV infectivity and death of SNaPs, we sought to identify natural genetic variants that affect the expression of these genes in SNaPs. We pooled 44 hESC-derived SNaP lines into a village (Village-44; Figure 4a) and three days later processed these cells for scRNA-seq and donor re-identification. As expected, cell cycle stage and differentiation status contributed to variation in gene expression (Figure 4b-c). We used scRNA-seq data to assess the quality and identity of the cells in this village, finding that SNaPs predominantly clustered with fetal NPCs when compared to an integrated reference dataset composed of human fetal and adult brain cells (Darmanis et al., 2015; Nowakowski et al., 2017) (Figure S5a). SNaPs and fetal NPCs expressed progenitor markers *HES1, SOX2*, and *VIM*, while displaying little to no expression of markers of differentiated neurons or astrocytes (Figure S5b). Furthermore, using the cell type classification pipeline in Seurat 3.0 (Stuart et al., 2019) to measure SNaP similarities to *in vivo* cells, we found that 83.1% of the SNaPs were determined to most closely resemble fetal NPCs, suggesting that the SNaPs have an expression profiles similar to those of their *in vivo* counterparts. In addition to confirming the relevance of this cell type to its *in vivo* counterpart, analysis confirmed the consistent production of high-quality SNaPs across all donors (Figure S5c-d).

**Figure 4:**
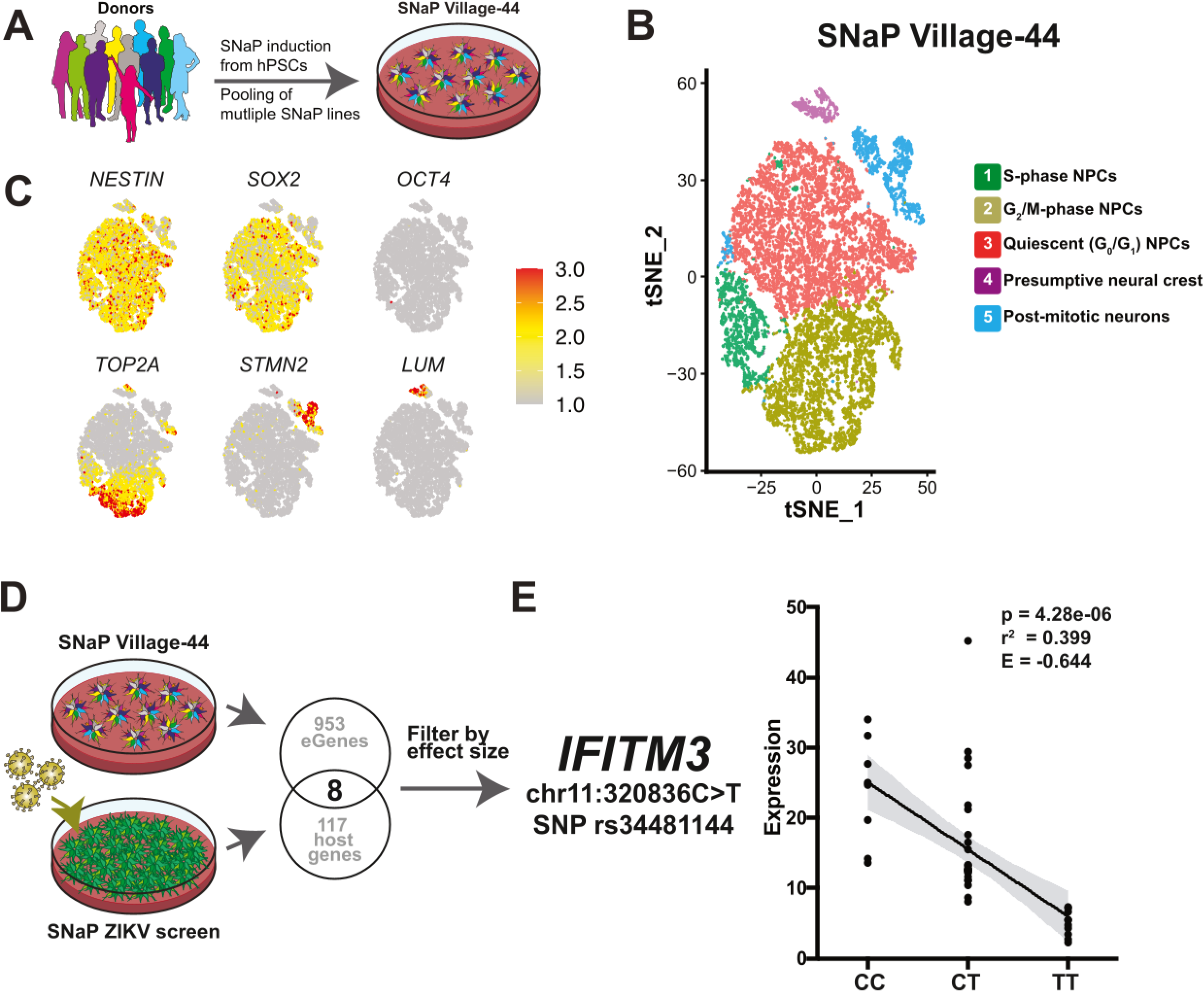
Differential expression of ZIKV host factors in SNaP villages. **A**, Schematic describing construction of 44-donor SNaP village. **B-C**, scRNA-seq of Village-44. (B) Though largely homogenous, SNaPs aggregate into five tSNE clusters driven primarily by cell cycle and differentiation status. (C) Expression of marker genes for NPCs *(NEST/N*, SOX2), pluripotent stem cells *(OCT4)*, cycling progenitors *(TOP2A)*, excitatory neurons (STM2) and presumptive neural crest *(LUM)*. **D**, Workflow combining Village-44 eQTL analysis with ZIKV survival screen results. 961 eGenes were filtered down to 8 potentially ZIKV-related genes with *IFITM3* having the largest effect size. **E,** *IFITM3* expression levels for SNaP lines in Village-44 harboring reference and alternate alleles at locus chr11:320836 (SNP rs34481144). Linear regression line in black with shaded error in gray.

We identified 961 genes whose expression levels were associated with the genotypes of nearby SNPs (“eGenes”, Data File 4). Many of these genes have known roles in fetal brain development. They included genes associated with neurodevelopmental disorders, including *MAPK3, PEX6*, and *DMPK* as well as the fetal brain transcriptional activator *CHURC1* and the Williams Syndrome genes *WBSCR16* and *WBSCR27* (Figure S5e, Data File 4). They also included many genes associated with autism spectrum disorder, epilepsy, and microcephaly such as *DPP10, HNRNPU*, and *TSEN2*. This analysis provides an inventory of genes regulated by common genetic variation in human NPCs, which could be useful for future investigations of variant effects on a range of neurodevelopmental processes and diseases.

We had hypothesized that screens of the human genome using synthetic (CRISPR) and natural (genetic) perturbations might converge. We cross-referenced the eGenes from our eQTL analysis to a list of 125 genes that were significantly enriched or depleted in both ZIKV survival screens. We found an overlap of 8 genes (Figure 4d) including the replication factors *EMC3* and *COG4*, and the antiviral genes *IFNAR1* and *IFITM3*. Of the 8 overlapping genes, SNPs in the *IFITM3* gene had the strongest effect size (i.e. the slope of the regression line normalized by mean expression of gene; E = -0.710 for rs6421983; Figure S5f). IFITM3 is believed to protect cells against RNA viruses by identifying and shuttling viral-containing endosomes to the lysosome for degradation (Spence et al., 2019). Previous work in immortalized cancer cell lines has shown that IFITM3 knockdown enhances ZIKV infection rates, while overexpression virtually eliminates viral replication (Brass et al., 2009; Savidis et al., 2016b). The influence of natural variation in *IFITM3* levels on flavivirus infectivity has yet to be explored.

We first aimed to pinpoint which *IFITM3* SNP was most likely responsible for the varied expression levels of this gene across cell lines. Pairwise linkage disequilibrium (LD) analysis of all statistically-significant *IFITM3* SNPs revealed four SNPs that were in high LD with the index SNP and thus associated to ZIKV infectivity with similarly large effect size (|E| > 0.6) (Figure S5g; Data File 4). Of these SNPs, rs34481144 (the only SNP located in a non-intronic region) is positioned in the 5’-UTR promoter segment of *IFITM3* and has previously been shown to influence expression of this gene (Allen et al., 2017), suggesting that it is the specific variant responsible for the effect we see. SNaPs from donors homozygous for the reference allele (C) of rs34481144 expressed *IFITM3* at levels 4.82-fold higher than donors homozygous for the alternate allele (T) (p = 4.28 × 10^−6^, r^2^ = 0.399; Figure 4e). We hypothesized that by enhancing expression of this antiviral gene the reference allele of rs34481144 confers protection from ZIKV infection, relative to the alternate allele.

### rs34481144 is a functional QTL influencing NPC sensitivity to ZIKV infection

To formally test the hypothesis that the reference allele protects SNaPs from ZIKV infection, we exposed SNaP Village-44 to ZIKV-Ug (MOI = 1) or mock media (Figure 5a). At 54 hpi, we FACS-sorted the cells into four fractions based on ZIKV envelope protein 4G2 antibody-stained GFP signal intensity (ZIKV-Negative, -Low, -Mid, -High) before harvesting pellets for DNA sequencing. We then analyzed each cell fraction using Census-Seq (Mitchell et al., 2020) to estimate each donor’s cellular contribution to the different fractions. Donors with the rs34481144^TT^ genotype were greatly over-represented relative to rs34481144^CC^ donors in the ZIKV-positive populations relative to the ZIKV-negative pool (r^2^ = 0.225, p = 3.04 × 10^−3^; Figure 5b-e). No such relationship was observed with other significant Village-44 eQTLs located in antiviral/host factor genes (Figure S6a). These data support the hypothesis that the rs34481144-T allele renders SNaPs more vulnerable to ZIKV infection compared to cells harboring the rs34481144-C allele (Figure 5f).

**Figure 5:**
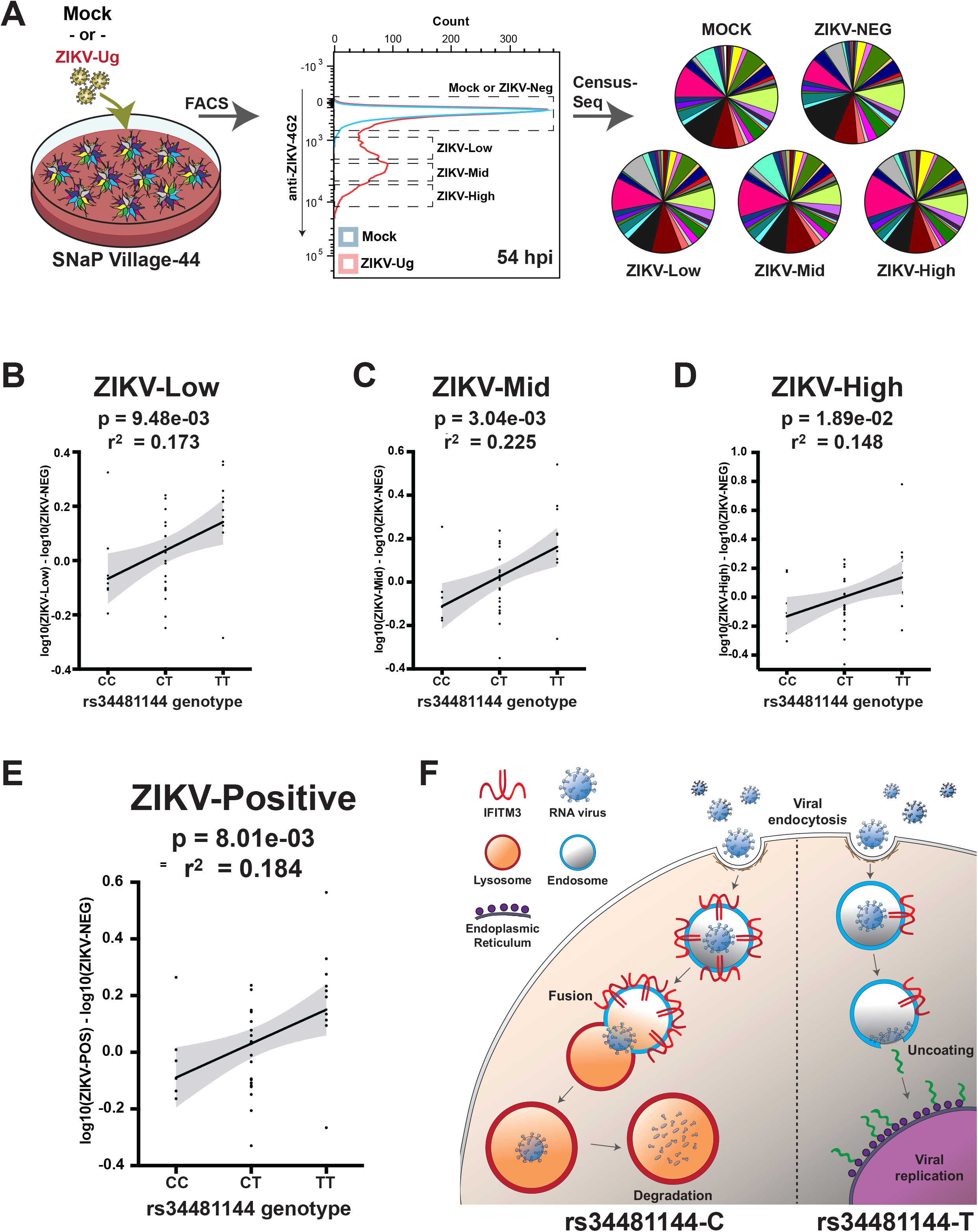
SNaP sensitivity to ZIKV associates to a common SNP in *IFITM3*. **A**, Experimental workflow. SNaP from 44 donors were cultured together, then infected with ZIKV-Ug (MOI = 1) or mock media. At 54 hpi, cells were FACS sorted into 4 bins based on 4G2 signal intensity. Representations of the individual donors in each of the FACS-sorted cell fractions was measured by Census-seq analysis, which infers the quantitative representation of each donor in a mixture from the quantitative representations of hundreds of thousands of common SNP alleles in that mixture. **B**-**E**, Association between genotype for rs34481144 and the distribution of different donors’ SNaPs between the ZIKV-positive fractions (ZIKV-Low, -Mid, -High, as well as an aggregated ZIKV-positive bin), relative to ZIKV-negative fraction. All values were normalized to mock controls to control for differences in intrinsic growth rate differences across donors in the village. Linear regression line in black with shaded error in gray. **F**, Schematic model, in which decreased expression levels of *IFITM3* (which associate with the rs34481144-T allele) result in increased ZIKV infectivity in SNaPs.

We next asked whether the apparently strong effect of the rs34481144 genotype on ZIKV susceptibility is cell-intrinsic or arises in an unexpected way from the cell-village experimental design. To do so, we analyzed data from SNaP infectivity assays of independently-cultured individual SNaP lines (Figure 3b) and again found a significant correlation between rs34481144 genotype and the percentage of ZIKV-infected SNaPs (at 54 hpi) in each culture, where the rs34481144^TT^ cells were more prone to infection than rs34481144^CC^ SNaPs (r^2^ = 0.345, p = 2.53 × 10^−3^; Figure 6a-b). Plotting these results against the Census-seq data showed agreement of results between the pooled-culture and arrayed-culture experimental designs (r^2^ = 0.703, p = 2.63 × 10^−5^; Figure S6b).

**Figure 6:**
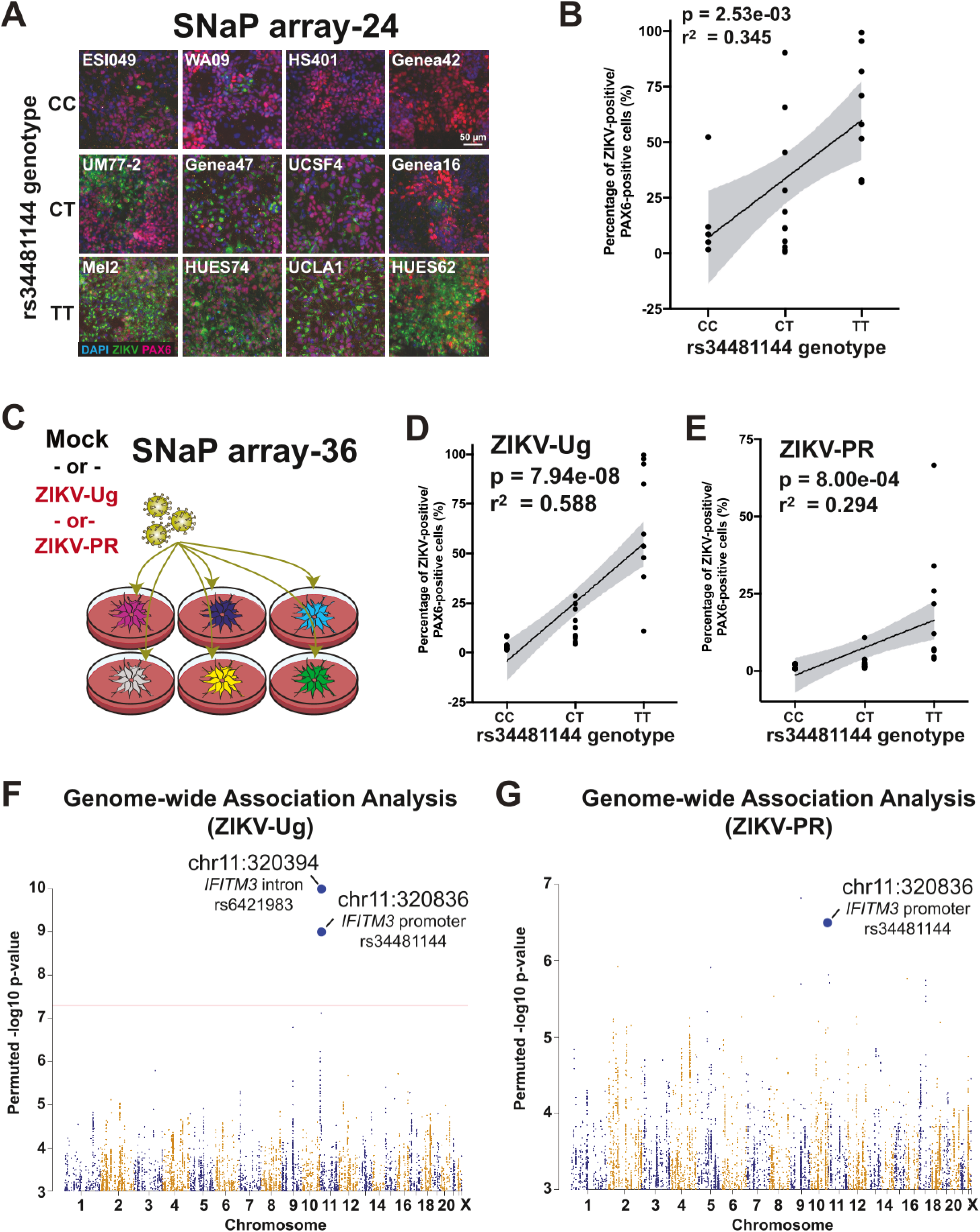
Further validation of the *IFITM3* SNP’s role in ZIKV infectivity. **A**, Representative images from 24 SNaP lines infected in arrayed format; rows correspond to donor’s rs34481144 genotype. **B**, Association of infectivity to rs34481144 genotype. **C**, Schematic showing arrayed infection of 36 SNaP lines with mock media, ZIKV-Ug (MOI = 1), or ZIKV-PR (MOI = 10). **D-E**, Quantification of (D) ZIKV-Ug and (E) ZIKV-PR arrayed infection phenotype to rs34481144 genotype. Linear regression line in black with shaded error in gray. **F-G**, Genome-wide association analysis of ZIKV infectivity across SNaPs from 36 donoros infected with (F) ZIKV-Ug and (G) ZIKV-PR. Red line denotes genome-wide significance.

To further replicate these results, we infected a separate set of 36 human iPSC-derived SNaP lines in an arrayed format using ZIKV-Ug (MOI = 1) or ZIKV-PR (MOI = 10). In these experiments, rs34481144 genotype explained 58.8% and 29.4% (respectively) of the variation in cell survival (Figure 6c-e).

Our focus on the rs34481144 SNP had been driven by the implication of this gene in our CRISPR-Cas9 and eQTL screens. We wondered whether this SNP’s effect on ZIKV infectivity was sufficiently strong to have been identifiable in an unbiased genome-wide search for effects of common variants on this cellular phenotype. We thus performed genome-wide association analysis on the survival data from the 36 arrayed cell lines, measuring the association of this phenotype with more than 1 million common SNPs. rs34481144 was the human genome’s top-scoring SNP in the ZIKV-Ug dataset, reaching genome-wide significance even in this modest 36-donor sample (p = 9.0 × 10^−10^; Figure 6f). In an analogous analysis of infectivity with the PR strain of ZIKV, rs34481144 also associated strongly, but just below genome-wide significance (p = 3.2 × 10^−7^; Figure 6g), and was the genome’s second strongest common-variant association.

These results demonstrate that a genetically variable, cell-intrinsic property of human NPCs is the major source of inter-individual variation in their susceptibility to a viral pathogen.

## DISCUSSION

Through analyses of natural and synthetic genetic perturbation on cellular phenotypes, we found that IFITM3 limited ZIKV infection of NPCs, and that a common SNP allele (rs34481144-T) in the *IFITM3* promoter accounted for 58% of the variation in ZIKV infectivity of human NPCs across dozens of genetic backgrounds. This same allele associated with three-fold reduction in expression of *IFITM3* in our experiments on NPCs from many donors (Figure 4e). Variants in the *IFITM3* gene, including SNPs rs12252 and rs34481144, have previously been associated with the severity of influenza infections (Allen et al., 2017; Pan et al., 2017). In PBMCs, the presence of the T allele at rs34481144 alters binding of transcription factors and reduces *IFITM3* promoter activity, leading to reduced baseline expression of *IFITM3* (Allen et al., 2017). Based on our findings, we believe that the rs34481144-T allele confers similar vulnerabilities to developing human brain cells exposed to ZIKV.

Intriguingly, frequencies of the rs34481144-T allele are highly variable across different global ancestries (Auton et al., 2015; Kim and Jeong, 2020), with frequencies as high as 46% in Europe and as low as 0.6% in regions in which flaviviruses are endemic. An intriguing possibility is that evolutionary selection has historically favored the rs34481144-C allele in places with high rates of mosquito-borne RNA virus infections.

This presents a concerning though likely scenario in which climate change-induced spread of mosquito vectors into Europe (Kraemer et al., 2019) introduces flaviviruses to populations that lack the natural defenses afforded by the protective *IFITM3* allele. Our results suggest that genotypic screening at the rs34481144 locus might be useful in identifying at-risk individuals during these predicted future outbreaks, and supports the enhancement of *IFITM3* expression levels as a potential therapeutic approach in the fight against this growing threat (Zou et al., 2021). Increases in *IFITM3* expression could potentially be accomplished through FDA- and EMA-approved Type I interferon treatments, some of which were recently determined to be safe for pregnant women (Hellwig et al., 2020).

A clear and timely extension of the kinds of analysis in this work could identify SNPs that affect rates of infection and death of human lung cells by SARS-CoV-2, the pathogen responsible for the global pandemic of coronavirus disease 2019 (COVID-19). Previous work has shown that IFITM3 activities can restrict SARS-CoV and MERS-CoV infections (Huang et al., 2011; Wrensch et al., 2014; Zhao et al., 2014), and a recent study in a relatively small number of patients found an association between rs12252 genotype and COVID-19 severity (Zhang et al., 2020), though this was not observed in a larger dataset (COVID-19 Initiative, 2021).

Outside the realm of human-viral interactions, the population-scale culturing of SNaPs could make it possible to map genetic underpinnings of variation in neurodevelopmental outcomes. Such SNaP villages could be exposed to effectors of major signaling pathways, such as WNT or Notch, could reveal the ways in which genetic profiles impact some of the earliest molecular events of cortical neurogenesis, while exposure to lead and other heavy metals could explain differential susceptibilities to environmental toxins across individuals. Population-scale analysis of SNaP villages could seek answers to basic questions about the etiology of neurodevelopmental disorders, such as the relationship between abnormal NPC proliferation and autism spectrum disorders (Deshpande et al., 2017; Marchetto et al., 2017; Qureshi et al., 2014), as well as the molecular and cellular mechanisms underlying genome-wide association results for autism spectrum disorders and connecting them to other traits, such as brain size.

Population-scale *in vitro* culture systems provide promising ways to capture the influence of genetic variation on cellular phenotypes (Jerber et al., 2021; Kang et al., 2018; Mitchell et al., 2020). We hope that these and many other new approaches open up new ways to find and characterize the many genetic and environmental factors that shape human development.

## ACKNOWLEDGEMENTS

This work was supported by NIH/NIMH grants U01MH105669 and U01MH115727, by the Stanley Center for Psychiatric Research at the Broad Institute and the Harvard University Faculty of Arts and Sciences Dean’s Competitive Fund for Promising Scholarship. M.F.W. is supported by the Burroughs Wellcome Fund Postdoctoral Enrichment Program award (1018707) and a K99/R00 Pathway to Independence Award (NIH/NIMH 1K99MH119327-01). We thank Janell Smith and Dr. Martin Berryer (Broad Institute) for their assistance in maintaining cell cultures. We acknowledge Olivia Bare, Adam Brown, and David Root (Broad Institute) for their contributions to the design and execution of the CRISPR-Cas9 screen.

## AUTHOR CONTRIBUTIONS

Conceptualization and design: MFW, JN, SG, JMM, FP, KE, SAM; Experimentation: MFW, JN, SG, JMM, CM, DM, KR, MT; hPSC line expansion, QC, and sequencing: MT, DH, supervised by RN. Dropulation code: JN, SAM; Data analysis: MFW, JN, SG, DM, DH, BKP; H1-Cas9 generation and validation: SK, KC, JJR, KAW; Writing: MFW, JN, SG, JMM, KE, SAM. Supervision: KE, SAM; Project administration: AN

---

**Supplemental Figure 1:**
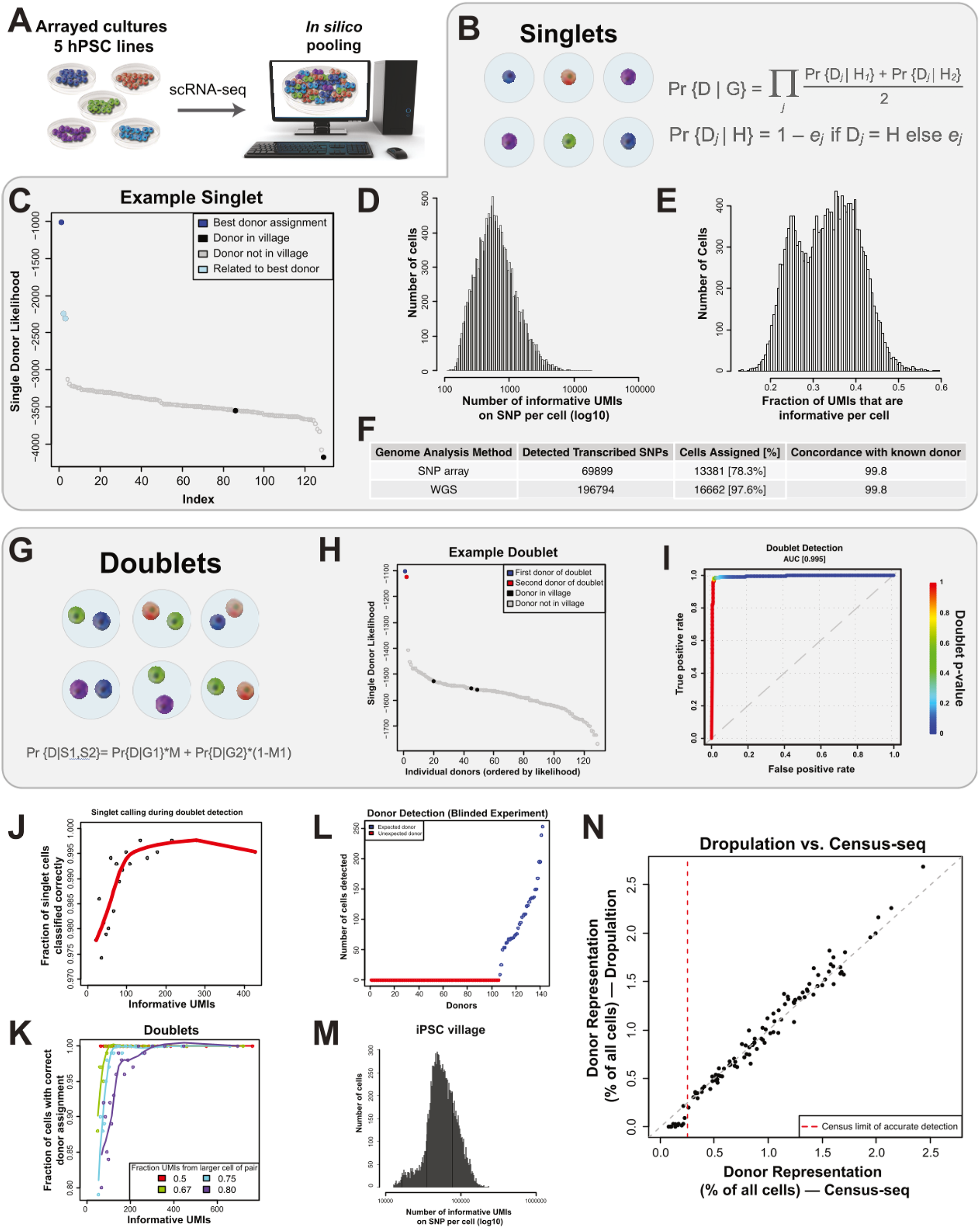
In silico evaluation and optimization of the analytical pipeline for cell-to-donor assignment (Dropulation). **A**, Schematic of *in silico* experiments. Five human pluripotent stem cell lines were cultured separately (i.e. in an arrayed format) prior to scRNA-seq preparations. Expression data was then pooled computationally for subsequent experiments. **B-F**, Dropulation singlet detection and donor re-identification pipeline. (B) Donor-assignment likelihood calculation, for singlet cells. (C) Plot depicting analysis of donor identity for an example cell. The sequenced cell was correctly assigned to the donor of origin (dark blue circle), distinguishing this donor from two related individuals (light blue circles). (D) Number and (E) fraction of unique molecular identifiers (UMIs) per cell that contain allelic information about at least one informative SNP that varies among the candidate donors. (F) Table depicting the outcomes of donor re-identification attempts when donor genotypes were obtained using whole genome sequencing or SNP arrays. The availability of more-complete genetic data (WGS) increases power in the analysis. **G-I**, Detection of cell-cell doublets. (G) Doublet detection likelihood function used. (H) Example cell in which two donors (red and dark blue circles) show similar likelihoods of assignment before doublets are considered. (I) Doublet detection algorithm successfully recognizes the presence of doublets for exclusion. Color along the line denotes p-value for the discrimination between best singlet and best doublet model. **J-K**, Requirements for successful singlet and doublet detection. Plot showing the number of informative UMIs needed for accurate (J) single cell donor assignment and (K) doublet detection. **L**, Results of blinded experiment identifying donors from a much-larger set of candidates. All 36 cell donors in this test hPSC village were detected and assigned to sequenced cells (blue dots). No cells from >100 other candidate donors (red dots) were detected. **M**, Number of UMIs-per-cell containing allelic information informative SNPs per cell in a 104-donor hiPSC village dataset. **N**, Concordance of estimates of hiPSC village donor composition obtained through Dropulation (by single-cell RNA-seq) and Census-seq (by bulk DNA-seq) analyses.

**Supplemental Figure 2:**
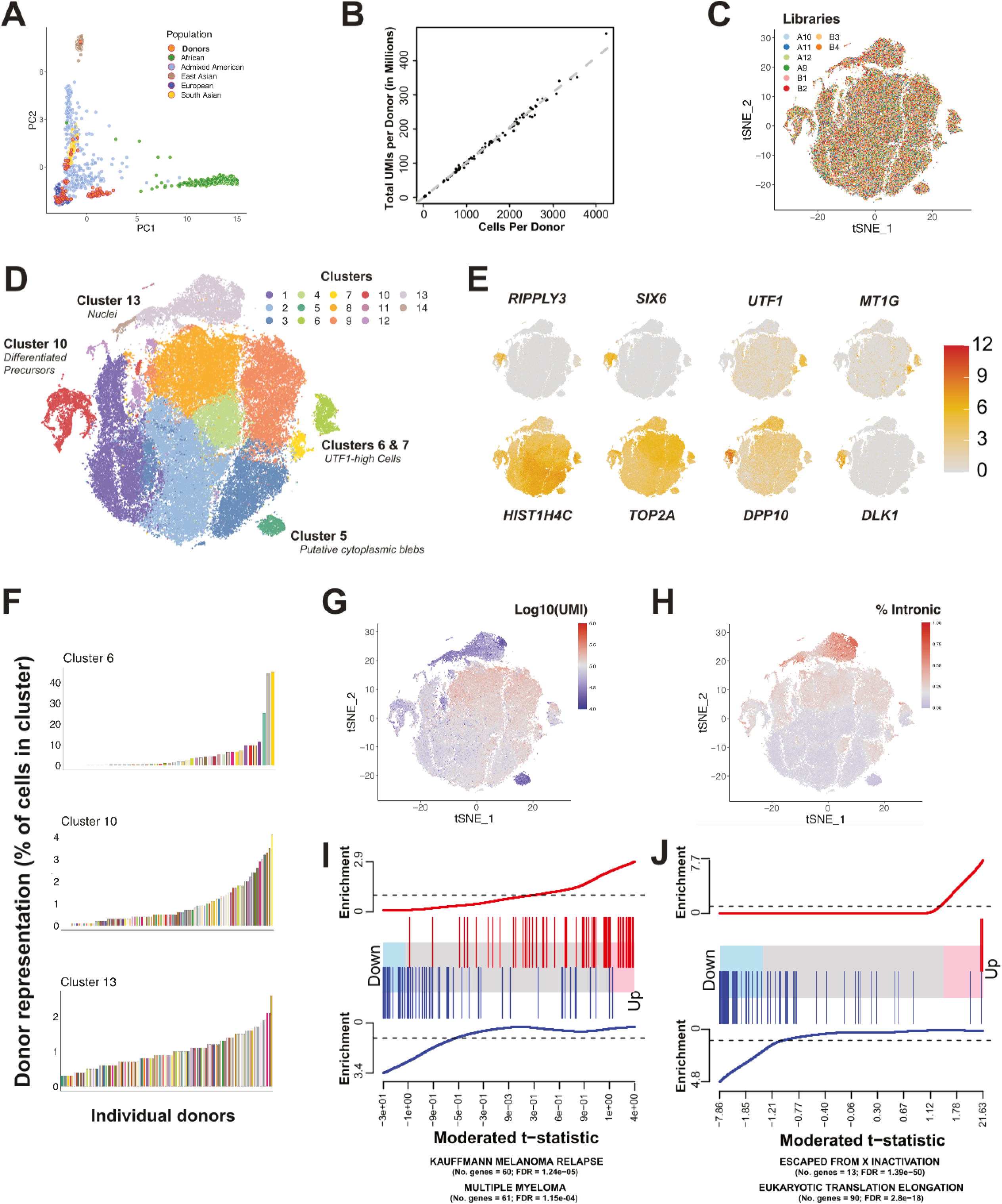
Sources of variation in 104-donor hiPSC village. **A**, PC plots depicting ancestral backgrounds of iPSC donors (orange circles). **B**, Total UMIs per donor for hiPSC village. **C**, tSNE plot of village scRNA-seq data color-coded by scRNA-seq 10X library. **D**, tSNE clusters from village scRNA-seq. Sequenced cells grouped into 14 unique clusters representing proliferative stem cells, differentiated cells, nuclei, and cytoplasmic blebs. **E**, Expression of marker genes for mitotic stem cells (*HIST1H4C, TOP2A*), differentiated neural cells (*RIPPLY3, SIX6, DLK1, DPP10*), and differentiation potential-associated genes (*UTF1, MT1G*). **F**, Breakdown of donor representation in three representative clusters. Three cell lines represented a majority of low differentiation potential (“UTF1-high”) cluster 6 cells. **G-H**, Cluster 13 cells are likely nuclear fragments as demonstrated by the (G) low numbers of UMIs and (H) high percentage of intronic reads. **I-J**, CAMERA gene enrichment plots highlight signatures detected from (I) cell source and (J) donor sex differential expression analysis.

**Supplemental Figure 3:**
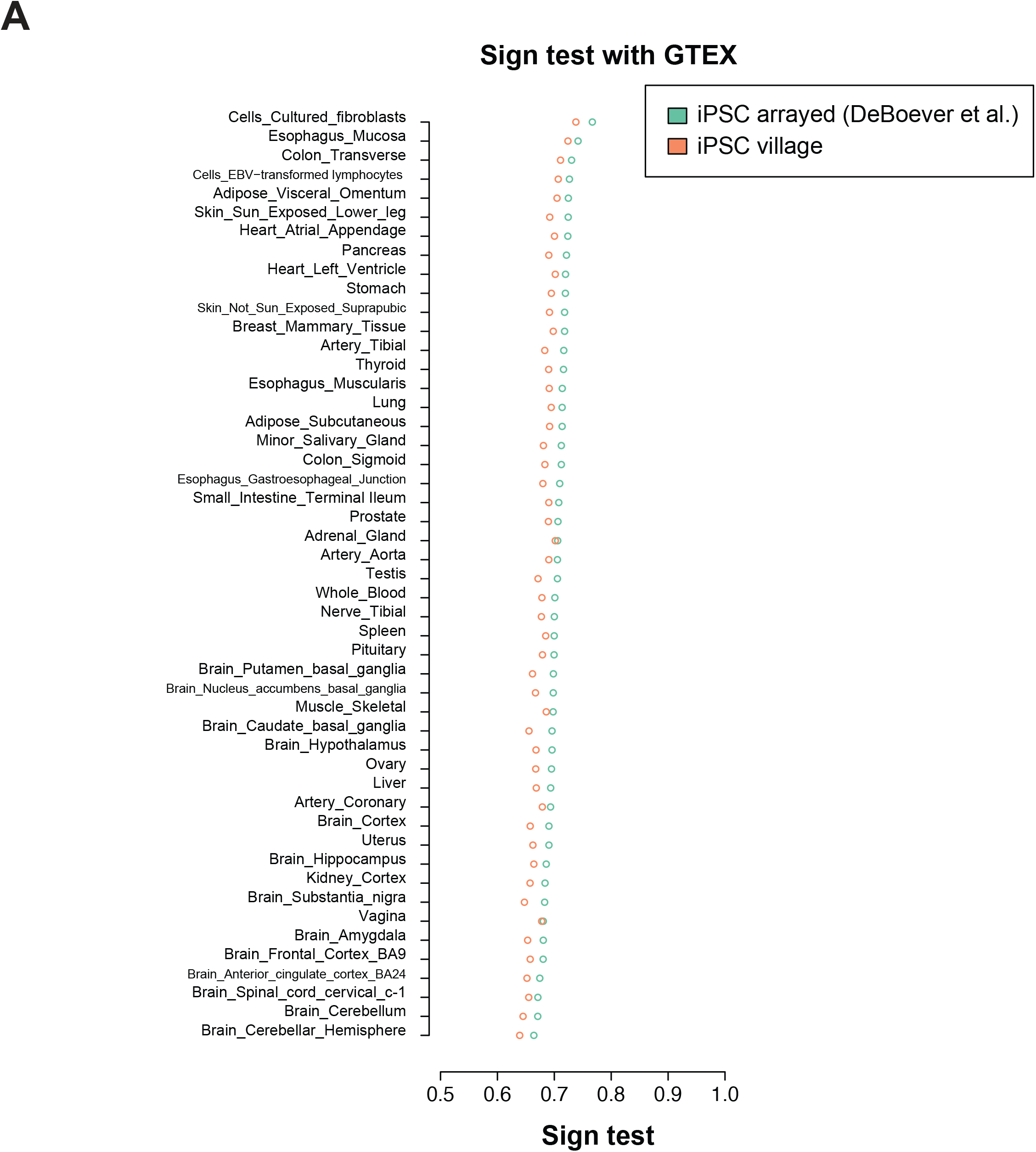
hiPSC village eQTL sign test comparison with human tissue dataset. **A**, Fraction of SNPs from arrayed and village iPSC datasets that showed the same direction of association to the gene-expression phenotypes in other studies. Similar sign test results were observed between the village and arrayed data when compared to GTEX tissue and
cell type data.

**Supplemental Figure 4:**
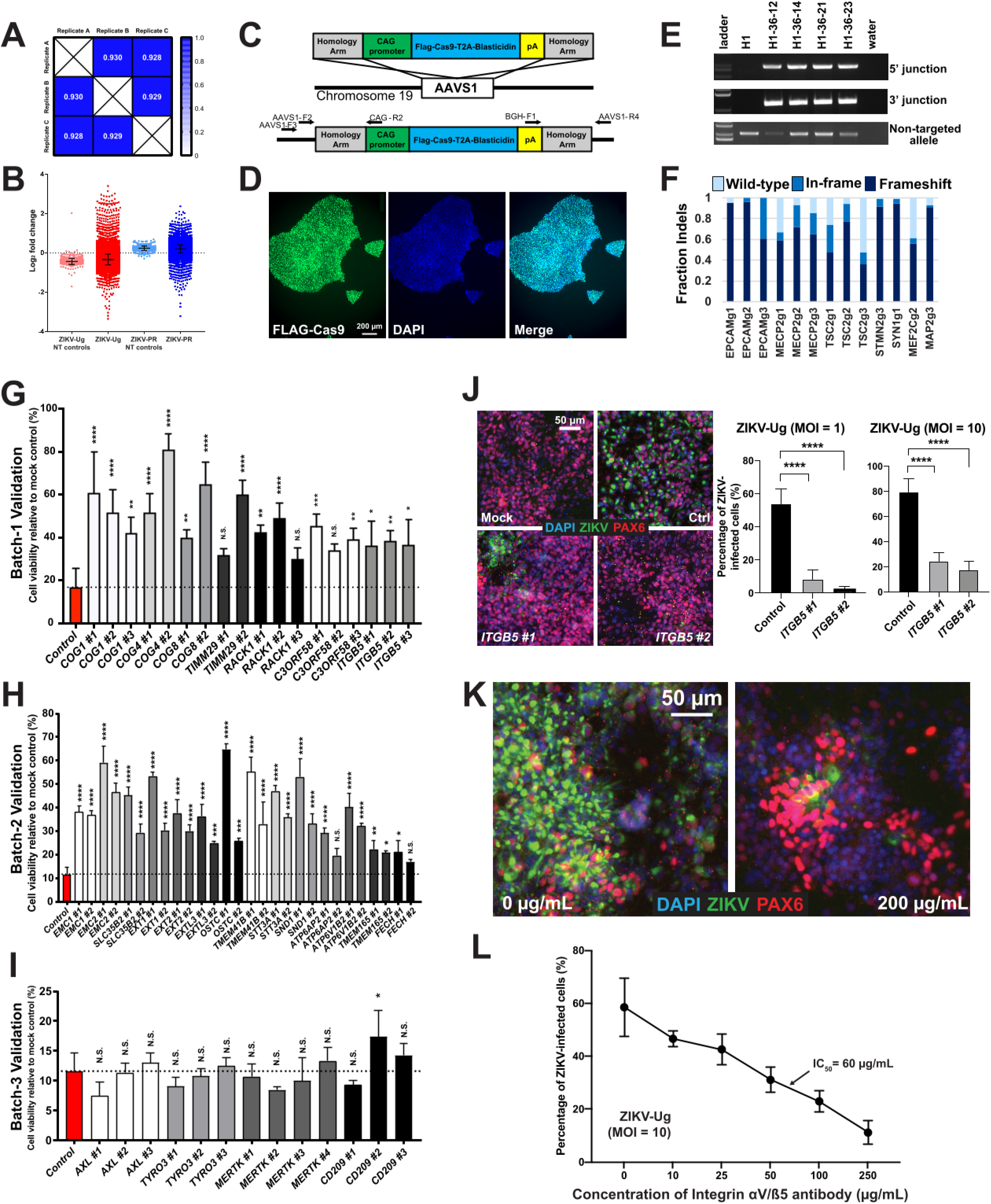
Quality controls and enrichment analysis results of SNaP ZIKV survival screen. **A**, Correlation matrix of Pearson r values across all replicates (range 0 to 1). **B**, Log2 fold change of individual guides in the ZIKV-Ug (red dots) and ZIKV-PR (blue dots) survival screens. Non-targeting (NT) controls (pink, light blue dots) are also shown. **C**-**F**, Validation of H1-Cas9 stem cell line. (C) Diagram of AAVS1 targeting vector and site of genomic integration. (D) Immunostaining using an anti-FLAG antibody for clone H1-36-23. Scale bar = 200 µM. (E) PCR across the junctions for 4 clones, indicating proper targeting into one allele. (F) Fraction indels in neurons differentiated from H1-36-23 clone and infected with lenti-gRNAs, measured by next-generation sequencing. **G**-**I**, Confirmation of primary screen hits. H1-Cas9 SNaPs were transduced with individual gRNAs, expanded, and exposed to ZIKV-Ug (MOI = 1). At 120hpi, cell viability was measured relative to mock controls. Dashed line denotes viability levels for non-targeting gRNA controls in each batch. **J-L**, Integrin aVb5 as a ZIKV entry factor. (J) Genetic ablation of *ITGB5* decreases ZIKV-Ug (MOI = 1-10) infectivity relative to controls at 54 hpi. Scale bar = 50 µM. (K-L) Anti-integrin aVb5 antibody blockade prevents ZIKV penetration in a dose-dependent manner (IC50 = 60 mg/mL). Scale bar = 50 µM. One-way ANOVA with Dunnett’s test for multiple comparisons (G-J, L) was used for statistical analysis. Data presented as mean ± S.D. N.S. = not significant, *p<0.05, **p<0.01, ***p<0.001, ****p<0.0001.

**Supplemental Figure 5:**
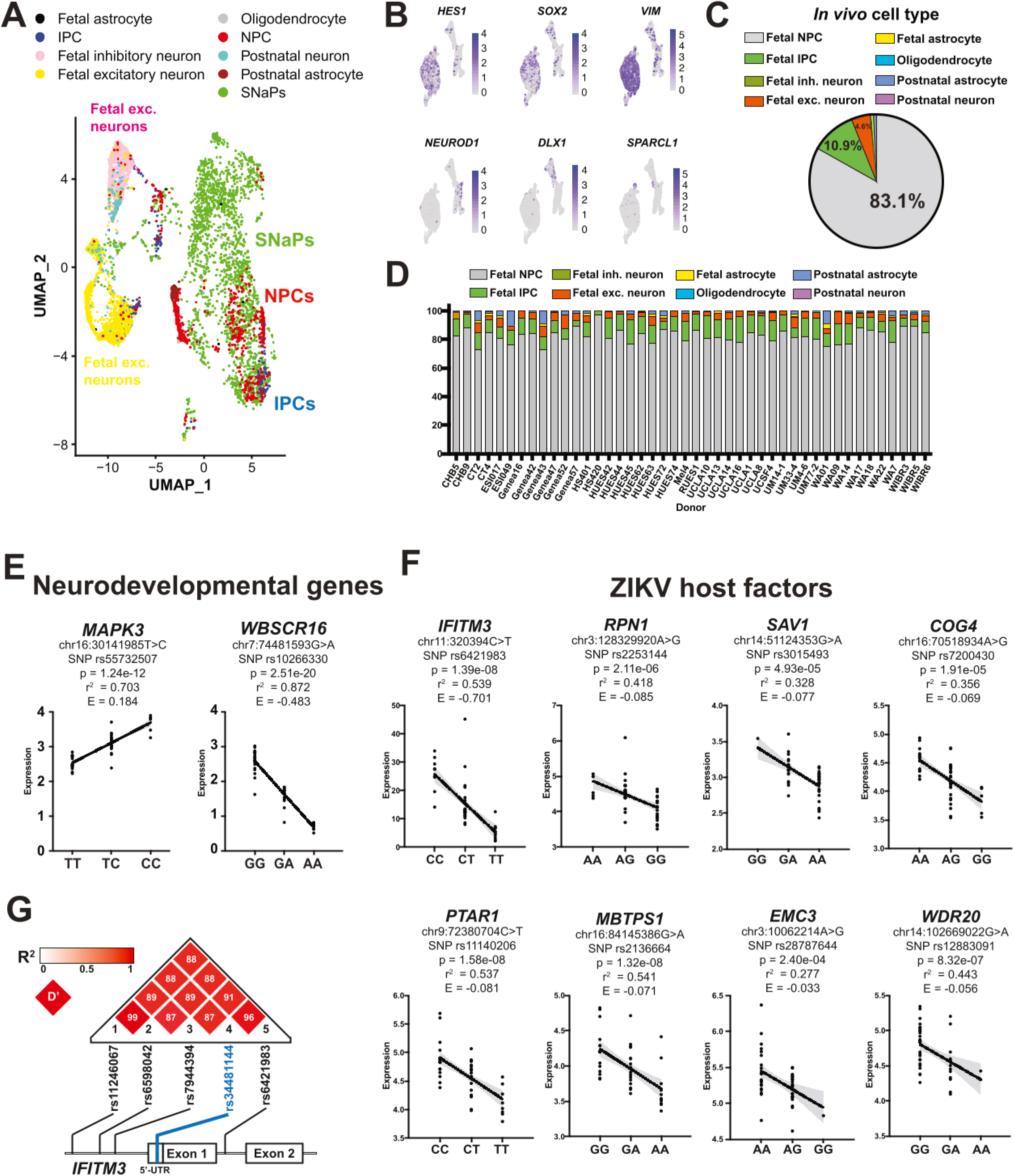
eQTL analysis reveals variation in gene expression across dozens of SNaP cell lines. **A**, UMAP cluster plot of Village-44 SNaPs and reference fetal and adult brain cells. **B**, NPC marker expression limited to SNaP/Fetal NPC cluster. Excitatory neuron (*NEUROD1*), inhibitory neuron (*DLX1*), and astrocyte (*SPARCL1*) markers are enriched in non-SNaP/Fetal NPC clusters. **C**-**D**, Quantification of computed cell type classification. (C) Seurat 3.0 computed cell type classification for all SNaPs in Village-44 and (D) on a per donor basis. **E**, Village-44 eQTL plots in genes relevant to human neurodevelopment. **F**, Village-44 eQTL plots in genes identified by the survival screens as ZIKV host or antiviral factors. Linear regression line in black with shaded error in gray. **G**, Linkage disequilibrium analysis of 5 statistically-significant eQTLs associated with *IFITM3* gene in Village-44. Box color depicts R^2^ value while number within square denotes D’ value of pairwise comparisons.

**Supplemental Figure 6:**
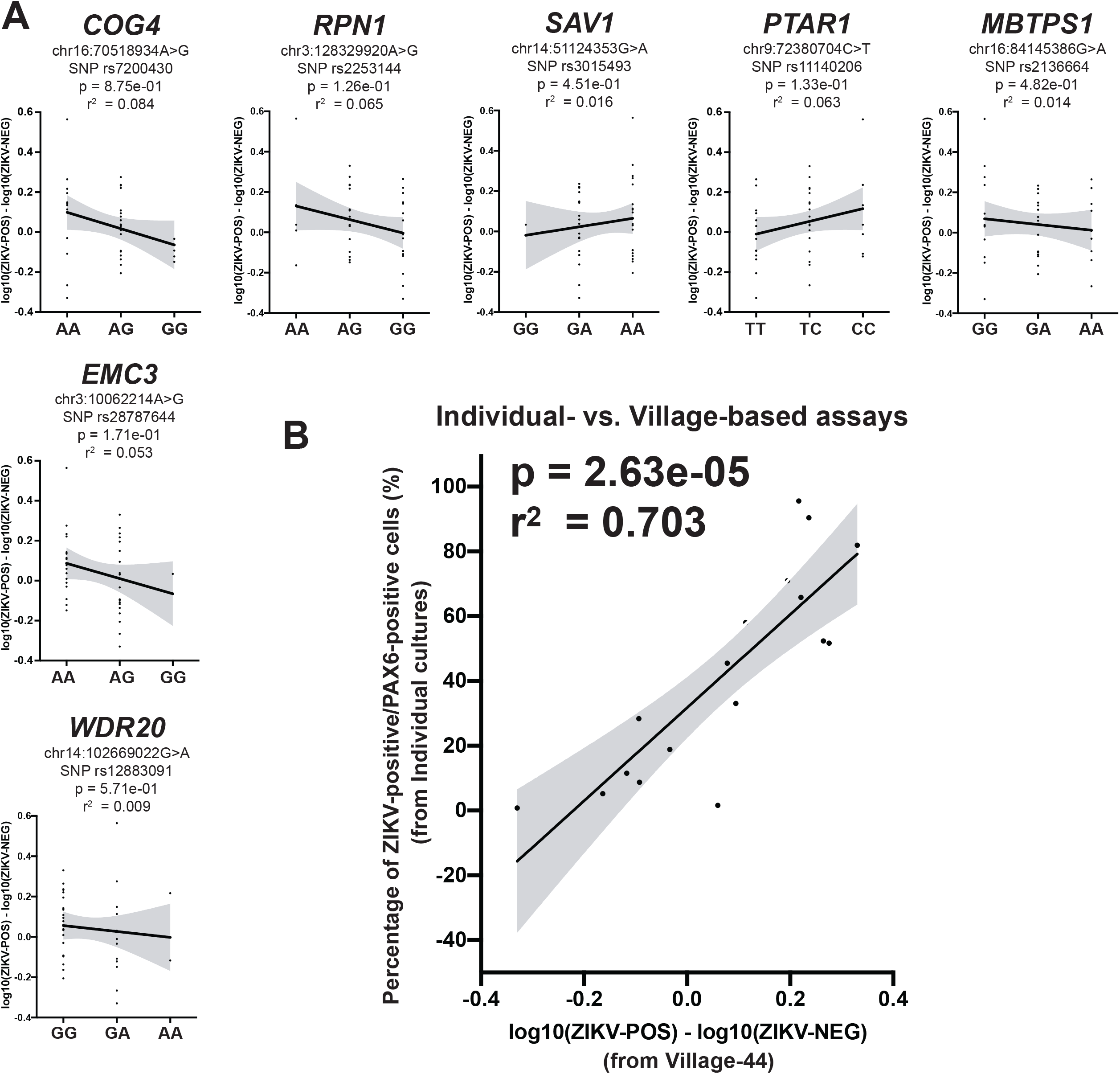
Natural variation in *IFITM3*, but not other antiviral genes, correlates with ZIKV infection severity. **A**, Village-44 eQTLs in ZIKV non-*IFITM3* host factor genes show no relationship with infectivity of SNaPs. **B**, Synergy between Village-44 Census-seq results and independently-cultured infectivity assay data. Linear regression line in black with shaded error in gray.

## SUPPLEMENTAL DATA FILES

Data File 1: Human iPSC village differential gene expression and CAMERA analysis

Data File 2: Human iPSC village eQTL results

Data File 3: ZIKV CRISPR screen results

Data File 4: SNaP village eQTL results

