## Supplemental Note for "Natural variation in gene expression and Zika virus susceptibility revealed by villages of neural progenitor cells"

**Supplementary Note:**
**Assignment of cells to individual donors using transcribed SNPs.**

To critically evaluate the accuracy with which cells were assigned to individual donors, we first cultured and analyzed (by scRNA-seq) lines from five distinct donors, then mixed the data *in silico* to create a “virtual cell village” for which the true donor of each cell was known (Figure S1a). We implemented a maximum likelihood approach for donor re-identification, in which for each cell, each observation of a transcript (an informative unique molecular identifier or UMI) carrying an allele (for a variable SNP) increased the likelihood of those donors who have that allele in their genomes, relative to other donors (Figure S1b). The change in donors’ likelihoods incorporated an uncertainty term, estimated from the data itself, to accommodate the possibility of sequencing error, genotyping error, or ambient/background RNA. By integrating observations across the hundreds of variant sites ascertained in a cell’s mRNAs, a single donor typically emerged as many orders of magnitude more likely to be the source of the cell than any other donor (**Figure S1c**). For these *in silico* mixing experiments, for which we knew the correct donor of each cell, we compared the predicted donor identities to the known ones. A calculation for an example individual cell is shown in Figure S1c: the known donor of origin of this cell is also the donor determined to have the highest computed likelihood; all other possible donors, including the two donors who were genetically related to the most likely donor, showed far-lower likelihoods for this cell (1,232.8 log10 decrease, 18,511 informative UMIs across 8083 SNPs). (Results like this indicate that donor assignment can confidently distinguish even among close relatives; it should be noted, however, that the approach would not be able to distinguish between monozygotic twins, nor between different cell lines from the same donor.)

It was important to evaluate the sensitivity and specificity of the analysis. scRNA-seq captures thousands of transcripts per cell, a subset of which contain transcribed SNPs. A SNP is informative if we know its genotype to vary among the cell donors, which we determined from whole genome sequencing (WGS) data we generated on the individual donors before the experiment. On average, single cells contained hundreds to thousands of informative transcripts (Figure S1d), representing 20-50% of the UMIs ascertained in the cell (Figure S1e). However, some individual cells with low RNA ascertainment did not have a sufficient number of transcripts to confidently re-identify the donor of origin; this is seen in that no donor greatly exceeded the other donors’ likelihood scores. By considering this distribution of assignment probabilities acros donors, we determine that these cells are instead “not confidently assigned” to any one donor. This number of unassigned cells was small (2.4%) when whole genome sequencing (WGS) data were available for the individual donors (as in this work), and larger when we limited analysis to SNPs from a common genotyping array (Illumina GSA); in both scenarios, however, the frequency of donor mis-assignment by the algorithm was low (0.2%) (Figure S1f).

A common challenge during scRNA-seq preparations involves “doublets”, which occur when a droplet encapsulates more than one cell, such that their cDNAs receive the same cell barcode and cannot be distinguished from one another. Detecting and removing doublets is an important challenge in single-cell analyses. Allelic information provides powerful ways to detect doublets in cell mixtures (Kang et al., 2017): since the great majority of doublets involve cells from two different donors, the resulting profile contains a mixture of alleles from two different donors. In this scenario, an *in silico* mixture of two donors' genotype data (Figure S1g) generates a higher likelihood (of the observed allelic observations in a single cell) then any one donor's genotype data does. To accommodate the possibility that the two cells will different amounts of RNA or informative UMIs, we add mixtures of a variety of proportions (4:1, 3:1, 2:1, and 1:1) to compete with the single-donor models.

When a doublet containing two cells from distinct donors at equal proportions (1:1) is analyzed by single donor assignment, both donors have similar probabilities of assignment (likelihoods for an example doublet are in Figure S1h). A doublet model, consisting of a 1:1 mixture of the two donors' genomes, generates the single cell's data with a far greater likelihood than either of the single-donor models does. In *in silico* evaluations, the doublet detection algorithm had a true positive rate of 98.3% and a false positive rate of 1.5% under these conditions, indicating that it could properly classify the great majority of doublets; misclassification of singlets as doublets was rare (Figure S1i). The number of informative UMIs in a cell is a critical factor for successful singlet and doublet calls. Singlets tended to be misassigned when they had fewer than 100 informative UMIs, as the false positive rate of cells with > 100 UMIs was 0.5% (Figure S1j). Doublets that were misclassified also had fewer informative UMIs. For ratios of 1:1, 2:1, and 3:1, 114 informative UMIs were sufficient to detect 99% of doublets correctly, and for 4:1 doublets, 238 informative UMIs were needed to reach this level of detection (Figure S1k).

Testing data with *in silico* mixing indicated that donors and doublets could be identified with high accuracy (99.8% and 98.95% respectively), but this analysis was not based on sequencing from donor lines cultured together as an *in vitro* village; when cells are grown and analyzed together, ambient or background RNA could be a substantial sounce of mis-assignment of alleles to cells. We therefore sought out to test these algorithms on data generated from a village of human cells grown together in a shared culture flask. We executed this experiment using two teams of researchers. One group ("wet lab") pooled 36 hESC donor lines into a village and processed scRNA-seq libraries for sequencing. The other group ("dry lab") received raw sequencing reads but was blind to the number, identity, and proportion of donor lines included in the village. After processing this data through the singlet and doublet detection pipelines, the dry lab group successfully identified all 36 donors (from a list of 142 candidates) present in the village without incorrectly assigning any cells to and of the 106 donors not included in the village (Figure S1l). The Dropulation software was able to analyze this dataset (25,080 cells sequenced to 1,755 average UMIs with 459M total reads) and assign cells to 142 possible donors in 1.72 hours using 512,235 SNPs, and detect doublets in an additional 1.4 hours, using 8g of memory on a single core processor.

To further test the ability of the Dropulation analytical pipeline to handle data generated in real-world experimental contexts, we constructed a village of human iPSCs from 104 distinct donors collected from the CIRM repository. Based on the number of informative UMIs per cell, the experiment was well powered to identify each donor of origin (Figure S1m). We then measured the relative proportion of each donor in the village using both Dropulation and Census-seq, a low-coverage-WGS-based computational tool we recently developed to infer the donor composition of cell mixtures from bulk DNA (without identifying the individual cells) (Mitchell et al., 2020). We observed a high level of concordance between these two methods for inferring donors' cellular representations in the mixture (for those donors present above the Census-seq detection limit of 0.3%), suggesting a high degree of concordance between methods that infer donor proportions in very different ways: from RNA and single-cell analysis (Dropulation), and from bulk DNA sequencing of the cell mixture (Census-seq) (Figure S1n). Together these findings validate the ability of the analytical approach to re-identify donors after the cells have been grown together in a common culture environment.
