## Supplemental Methods for "Natural variation in gene expression and Zika virus susceptibility revealed by villages of neural progenitor cells"

### SUPPLEMENTAL EXPERIMENTAL PROCEDURES

*Stem cell culture*: Human ESCs and iPSCs were maintained in mTeSR media (Stem Cell Technologies, 85850) on Geltrex basement membrane matrix (1:100; Life Technologies, A1413301). Cells were split every 4-5 days (when they reached 80-90% confluency) using a 15 minute/37°C incubation in Accutase (Innovative Cell Technologies, AT104) followed by 1:10 dilution in mTeSR. For each passage, media was supplemented with ROCK inhibitor Y-27632 (10 mM; Stemgent, 04-0012) for 24 hours after plating.

*Viral transduction:* TetO-Ngn2-Puromycin and Ubq-rtTA constructs were obtained from the Wernig lab (Stanford) before being packaged as high-titer lentiviruses (Alstem, Richmond, CA). When hPSCs reached 80-100% confluency, they were dissociated with Accutase before being re-suspended in lentivirus-containing mTeSR media supplemented with Y-27632 at a range of MOI = 1 to MOI = 3. Cells were then plated on Geltrex-coated 12-well plates at 500,000 cells per well in a total volume of 750 ml/well. After 18-24 hours, lentiviral media was aspirated and cells were fed with mTeSR media and maintained as described above. Transduced cells were maintained for up to 10 passages for inductions and transduction efficiencies typically ranged from 65-85% across cell lines.

*Induction of SNaPs from human PSCs*: Human PSCs were dissociated and plated at 75,000 cells/cm$^2$ on Geltrex matrix in mTeSR media supplemented with Y-27632. After 12-24 hours, cells were fed with Induction Media (Day 1): DMEM/F12 (ThermoFisher, 11320082), Glutamax (1:100; ThermoFisher, 10565018), 20% Glucose (1.5% v/v), N2 Supplement (1:100, ThermoFisher, 17502048), Doxycycline (2 mg/mL; Sigma-Aldrich, D9891), LDN-193189 (200 nM; Stemgent, 04-0074), SB431542 (10 mM; Tocris, 1614), and XAV939 (2 mM; Stemgent, 04-00046). After 24 hours in Induction Media, cells were fed with Selection Media (Day 2): DMEM/F12, Glutamax (1:100), 20% Glucose (1.5% v/v), N2 Supplement (1:100), Doxycycline (2 mg/mL), puromycin (5 mg/mL; ThermoFisher, A1113803), LDN-193189 (100 nM), SB431542 (5 mM), and XAV939 (1 mM). After 24 hours in Selection Media, SNaPs were dissociated with Accutase and replated at 120,000 cells/cm$^2$ on Geltrex-coated plates in SNaP maintenance media supplemented with puromycin and Y-27632 (Day 3): DMEM/F12, Glutamax (1:100), MEM-NEAA (1:100; Life Technologies, 10370088), B27 minus Vitamin A (1:50; Life Technologies, 12587010), N2 Supplement (1:100; Life Technologies, 17502048), recombinant human EGF (10 ng/mL; R&D Systems, 236-EG-200), recombinant human basic FGF (10 ng/mL; Life Technologies, 13256029), puromycin (5 mg/mL), and Y-27632 (10 mM). Starting 12-24 hours after passaging, SNaPs were fed daily with SNaP maintenance media lacking Y-27632 and puromycin. SNaPs were passaged every 5-7 days. All SNaP passages throughout this manuscript included overnight Y-27632 treatment.

*Village construction and experimental design*: Human iPSC and SNaP lines were maintained as independent cultures. At Passage 2-3, cell lines were dissociated with Accutase and counted using a Scepter Handheld Automated Cell Counter (Millipore Sigma, PHCC20060). An equal number of cells from each line were then plated together in a 10 cm$^2$ Geltrex-coated dish at 120,000 cells/cm$^2$. For Dropulation experiments, cells were harvested 2-3 days post-plating and run through the 10X Chromium Single Cell 3' Reagents V3 system to isolate individual cells into droplets per vendor's instructions (10X Genomics; San Francisco, CA). Samples were then sequenced on a NovaSeq 6000 system (Illumina) using a S2 flow cell at 2 x 100bp.

*Dropulation sequence alignment and donor assignment:* Raw sequence data was demultiplexed and aligned following the standard Drop-Seq protocol (Macosko et al., 2015). Human experiments were aligned to the GRCh38 reference and ensembl v89 gene models. Sequencing reads were then filtered to reads that mapped at high quality (MQ>=10) to the human genome. Genotype data was preprocessed as previously described (Mitchell et al., 2020). Genotypes in VCF files were called against the GRCh38 reference genome.

For Dropulation to perform accurately, input sequencing and VCF data is filtered on a per-run basis. Sequence reads are filtered to high quality mappings (MQ>=10) on the autosomes that have not been flagged as PCR duplicates. VCF sites are considered if they meet all of the following criteria: each site passed GATK's Variant Quality Score

Calibration (VQSR) filter, had a mean genotype quality (GQ) score ≥ 20, a mean variant read depth (DP) ≥ 10, a call rate > 50%, a Hardy Weinberg Equilibrium p-value > $1e^{-3}$. Variants located in low complexity regions of the genome or in common segmental duplications as annotated by the UCSC genome browser were filtered from the VCF. Individual genotypes with gross allelic imbalances were set to missing and excluded as defined by the following criteria: an allele balance ≥ 0.25 for heterozygous sites and ≥ 0.9 for homozygous reference and homozygous alternate sites. Samples with a call rate <90% and a mean depth < 10 were removed from the VCF. Additionally the Dropulation algorithm retains sites that: a) have a GQ score of at least 30 b) are diploid and polymorphic in the subset of donors in the population c) at least 50% of donors have a GQ score ≥ 30. Furthermore variants on the X, Y, and MT contigs were ignored. For genotype array based data where site quality scores may not be available, sites where the reference base is ambiguous [A/T, C/G] were not considered.

The Dropulation algorithm analyzes each cell in the data set independently, and generates a likelihood of the data having been generated by each of the donors in the VCF (or a subset of them as requested.) At each variant site, the probability of observing the allele at each unique molecular identifier (UMI) for the site is calculated as the probability of the base at that site for the mode observed base, and 1 - probability for reads that disagree with the mode UMI base. This downweighs transcripts where the underlying reads disagree on the observed allele. The likelihood of donor is then computed as the diploid likelihood at each UMI, summed across all sites. The diploid genotype is the average of the two haploid genotypes. For homozygous genotypes, this is the same as the haploid genotype, where if the observed base matches the genotype of the donor, the likelihood is 1- error rate of the UMI. For heterozygous sites, the likelihood is 0.5, regardless of base quality. The probability of the donor is then calculated as the probability of the donor divided by the sum of all donor probabilities.

*Dropulation missing data handling:* As the number of donors in a VCF file increases, the likelihood that at least one donor will not be called at high quality at any genotyping site increases. One way to deal with missing data is to ignore sites with any missing data, but this can exclude a large number of sites. Instead, we filter sites where the majority of donors are missing data, then for other sites missing data we use the remaining members of the population to calculate a per-site likelihood penalty score to use for all donors that have no genotype data. This score is an extension of the donor assignment score, where the likelihood of each genotype class is calculated, then combined as a weighted average score. This replaces the diploid genotype score for each UMI observed. The mixture coefficient is the proportion of the population that has each genotype class in the population, and sums to one.

The diploid likelihood for a single variant site:

$$\Pr\{D|G\} = \prod_j \frac{\Pr\{D_j|H_1\} + \Pr\{D_j|H_2\}}{2}$$

The haploid likelihood

$$\Pr\{D_j|H\} = 1 - e_j \text{ if } D_j = H \text{ else } e_j$$

The missing data penalty for a single UMI

$$PS = \sum_{i \in \{AA,AB,BB\}} Pr\{D|G_i\} \times M(G_i)$$

D=The list of UMI bases at the site
G=The genotype of the donor at that site.
H1,H2=The haploid genotype
ej= The error rate of the observed UMI at base Dj.
PS=The penalty score for missing data at a site
M=mixture coefficient [proportion of genotype in population]

*Dropulation doublet detection:* Doublet Detection uses the same read and variant filtering as the donor assignment algorithm, with the exception of missing data, where only sites with at least 90% complete data are accepted. The Doublet detection algorithm analyzes each cell in the data set independently, and generates a likelihood of the data having been generated by each possible pair of donors. To limit the number of possible tests, doublet detection is more restricted than donor assignment. The first donor of the pair is fixed as the most likely donor based on the single donor assignment, and the second donor of the pair is limited to a set of donors expected in the experiment. This limits the number of combinations to [number of donors -1] tests per cell.

For each donor pair, we optimize the mixture component of donor 1 to donor 2 to maximize the likelihood of that donor pair. The mixture score is the fraction of the data that arises from the first donor of the pair and is bounded to [0.8-0.2]. If the mixture score is unbounded, sequencing errors, ambient RNA, and genotyping errors will almost always generate mixtures of two donors that are very close to one, with a higher likelihood than the single donor likelihood, resulting in most cells being classified as doublets.

To select the donor pair that best explains the data, we first calculate the maximum likelihood each donor pair by selecting the maximum likelihood of the optimal mixture, the likelihood of the pair with a mixture of 1 (all data arises from donor 1) and the likelihood of the pair with a mixture of 0 (all data arises from donor 2.) The donor pair with maximum likelihood is then selected as the best pair.

To classify the pair as a singlet or doublet, we calculate the probability of the data being a doublet as the doublet likelihood divided by the sum of the doublet likelihood and mixture=1,0 likelihoods. We classify cells as doublets if their probability is greater >= 0.9. The vast majority of doublet probabilities are bimodally distributed at approximately 1 and 0.

Doublet Likelihood

$$Pr\{D|S_1, S_2\} = Pr\{D|G_1\} * M + Pr\{D|G_2\} * (1 - M)$$

$S_1$,$S_2$= Donor 1, Donor 2
$G_1$,$G_2$=Genotype Donor 1, Genotype Donor 2
M=Fraction of data arising from donor 1

*Dropulation assignable single donors*: To perform downstream analysis at a donor level, the set of cell barcodes in the experiment need to be filtered to the subset where cells are assigned to a single donor confidently. Cell barcodes are assigned to donors via single donor assignment. Doublet detection is then run, and all cells that are likely doublets (p-value > 0.9) are filtered from the data set. Cell barcodes are filtered if the single donor assignment p-value > 0.05. Given the remaining cells, the relative proportions of each donor are validated to determine if there are significant numbers of cells assigned to donors in the genotype backbone, but not expected in the experiment. Donors with very few assigned cells (less than 0.2%) are removed from the experiment.

*Single cell expression analysis of iPSC village*: The digital gene expression matrices were normalized and variable gene selection was performed as previously described (Saunders et al., 2021). Clusters were identified using Independent component analysis (ICA) based dimension reduction and Louvain community detection algorithms (Krienen et al., 2020).

*Differential gene expression and Geneset enrichment analysis*: To detect sex-biased or cell source dependent gene expression, we summed the UMI counts of all assignable single cells per donor to generate a donor by gene matrix. Differential expression was run as described in (GTEX paper) using voom-limma while adjusting for covariates including age and cell source where applicable and additional surrogate variables determined by smartSVA. Gene set enrichment was performed using the C2 (literature curated) and C5 (Gene Ontology Annotations) available in the Molecular Signatures Database (MSigDB) and CAMERA (Wu et al., 2012) on the list of genes ranked by the voom t-statistics.

*eQTL discovery*: For the set of assignable single donor cells, the UMI counts across cells of the same donor and gene are summed to a single measurement to generate a donor by gene expression matrix. Genes on the Y and MT chromosomes are filtered out, as are gene symbols that have ambiguous genomic mappings. Gene expression is then normalized to be fractional by dividing the gene expression of each gene/donor to the sum of expression for all genes. This fractional representation is then multiplied by a fixed constant of 100,000. Finally, genes are filtered to the top 50% highest expressed genes for eQTL discovery.

Variants are included for analysis if they pass all of the following filters. At least 90% of donors must have a genotype that was called and has a genotype quality >=30. The minor allele frequency of the variant must be between 5% and 95% in the population. The variant Hardy-Weinberg equilibrium (HWE) p-value must be > 1e-4. Finally, the variant must be within 10kb of the start or end of the gene for which it is tested. Variant genotypes are encoded by the number of alternate alleles.

The matrix of normalized expression data and genotype matrix per donor is then encoded in the format required by the R package MatrixEQTL (Shabalin, 2012). The expression data was then corrected for latent batch effects using PEER (Stegle, 2012). MatrixEQTL was then used on the corrected expression data to generate empiric p-values for all variant/gene interactions. False discovery rate (FDR) is then controlled hierarchically at two levels. At the gene level, the SNP with the best p-value is selected as the index SNP, and FDR is controlled by using the R package eigenMT (Davis et al., 2016), which uses the linkage disequilibrium of SNPs to determine the number of independent tests within a gene. The distribution of index SNP p-values is then transformed into q-values via the R package qvalue. We consider all genes with a q-value < 0.05 to be eGenes.

*SNaP cell type classification*: scRNA-seq based comparisons between the *in vitro* SNaPs and *in vivo* human brain cells were conducted using the Seurat 3.0 R package (Stuart et al., 2019). First, a custom script was composed based on the "Multiple Dataset Integration and Label Transfer: Reference-based" vignette (http://sajitalab.org/seurat/v3.1/integration.html; accessed July 15, 2019). Then, gene expression matrices from 13,732 Village-44 SNaPs were merged with two reference datasets: 257 cells from 16-18 week post-conception (wpc) fetal and 21- to 63-year-old adult brain tissue (Darmanis et al., 2015) and 3396 cells from 5.85 wpc to 37 wpc fetal brain tissue (Nowakowski et al., 2017). Similar to a recent report (Velasco et al., 2019), we condensed the large number of cell types identified in these reference datasets into 8 groups: Fetal Astrocyte (Nowakowski: "Astrocyte"), Fetal

Excitatory Neuron (Nowakowski: "EN-PFC-1", "EN-PFC-2", "EN-PFC-3", "EN-V1-1", "EN-V1-2", EN-V1-3", "nEN-early-1", "nEN-early-2", "nEN-late"), Fetal Inhibitory Neuron (Nowakowski: "nIN-1", "nIN-2", "nIN-3", "nIN-4", "nIN-5", "IN-CTX-CGE-1", "IN-CTX-CGE-2", "IN-CTX-MGE-1", "IN-CTX-MGE-2"), Intermediate Progenitor Cell (Nowakowski: "IPC-div1", "IPC-div2", "MGE-IPC-1", "MGE-IPC-2", "MGE-IPC-3", "IPC-nEN-1", "IPC-nEN-2", "IPC-nEN-3"), Neural Progenitor Cells (Darmanis: "fetal replicating"; Nowakowski: "RG-div1", "RG-div2", "RG-early", "vRG", "tRG", "oRG", "MGE-RG-1", "MGE-RG-2", "MGE-div"), Oligodendrocytes (Darmanis: "oligodendrocytes"), Postnatal Astrocyte (Darmanis: "astrocytes"), and Postnatal Neuron (Darmanis: "neurons").

After inputting the merged gene expression matrices with the condensed cell identifier metadata, the merged dataset was split into a list with each dataset as an element using the CreateSeuratObject function (min.cells = 3). Log-normalization was then performed and variable features were identified using the NormalizeData and FindVariableFeatures (selection.method = "vst"; nfeatures = 2000) functions, respectively. Anchors between the individual datasets were calculated using the FindIntegrationAnchors (dims = 1:30). The IntegrateData function (dims = 1:30) was deployed to create a batch-corrected expression matrix for all reference cells. This integrated expression matrix was then used as the reference to which the SNaP expression matrices ("query") were compared using the FindTransferAnchors, TransferData, and AddMetaData functions (dims = 1:30). The output included cell predictions and prediction scores for each SNaP cell.

*ZIKV infectivity assay:* SNaPs were infected for 1 hour at 37°C/5% $CO_2$ with ZIKV-Ug or ZIKV-PR diluted in 1X EBSS at a MOI of 10, 1, 0.1, or 0.01. Vero cell conditioned media from uninfected cells was diluted in 1x EBSS and used for the mock controls. Mock and ZIKV-infected cells were fixed 54 hours post-infection (hpi) with 4% paraformaldehyde for 15 minutes at room temperature and then washed with 1X PBS. Cells were permeabilized with 0.1% Triton for 15 minutes and then blocked with 10% normal donkey serum diluted in 1X PBS for 1 hour at room temperature followed by an overnight 4°C incubation in primary antibody: Mouse monoclonal D1-4G2 anti-flavivirus envelope protein (1:500) and Rabbit anti-PAX6 (1:500; Stem Cell Technologies, 60094) antibody diluted in blocking solution. After 3 washes in 1X PBS at room temperature, cells were incubated for 2-4 hours at room temperature in secondary antibody: Donkey anti-Mouse Alexa647 (1:1000) and Donkey anti-Rabbit Alexa555 (1:1000). Cells were washed once with 1X PBS followed by a 5-minute incubation in DAPI (1:5000). For Vero cell infections, an additional 20-minute room temperature incubation with F-Actin CytoPainter Phalloidin-iFluor 555 Reagent (1:10000; Abcam, ab176756) was included. Finally, cells were washed twice more with 1X PBS prior to imaging. For each well of a 96-well plate, 4-8 fluorescent images were taken using the Cytation 3 cell imaging multi-mode reader. All images were then processed using the CellProfiler imaging analysis software to quantify the percentage of 4G2-positive PAX6 stained cells.

*Genome-wide CRISPR-Cas9 ZIKV survival screens:* All gRNA and lentiviral reagents for the primary and validation screens were generated at Broad Institute Genetic Perturbation Platform. SW7338.1 SNaPs were generated and transduced with the Brunello barcoded sgRNA library (CP0043 Brunello library containing 77,441 barcoded sgRNAs targeting 19,114 genes and 1,000 not-targeting guides) delivered through the all-in-one LentiCRISPRv2.0 system (pXPR_BRD023 vector) (Doench et al., 2016; Sanjana et al., 2014). One hundred million SNaPs per replicate (3 total replicates) were transduced using the spinoculation method, in which cells were cultured in suspension with LentiCRISPRv2.0 (estimated MOI = 0.4) and centrifuged at room temperature for 2 hours at 1,000 rpm before being plated at 120,000 cells/$cm^2$ on Geltrex coated plates. Transduced SNaPs were then expanded and selected with puromycin (1 μg/mL) for one week, at which point they were passaged onto 15 $cm^2$ Geltrex-coated dishes at 120,000 cells/$cm^2$ (40 million cells were plated per replicate to maintain the 500 cells per sgRNA representation). Two days post-plating (one day post-Y27632 removal), SNaPs were either: (1) harvested using Accutase followed by PBS washes ("Pre-infection/Day 0" samples), (2) infected with mock media (for "Mock" samples), (3) infected with ZIKV-Ug (MOI = 1), or (4) infected with ZIKV-PR (MOI = 5) in minimal media for 1 hour at 37°C/5% $CO_2$ with gentle rocking every 15 minutes to prevent the cells from drying. Cells were then fed every other day starting at 48 hpi by removing all media, washing once with 1X PBS to remove dead cells and debris, and then adding back a 50:50 fresh SNaP maintenance media/conditioned media mixture. On Day 10, all samples ("Mock/Day 10", "ZIKV-Ug", and "ZIKV-PR") were harvested and frozen at -80°C. DNA was then extracted using the QIAmp DNA Blood Maxi kit (Qiagen, 51192). PCR and sequencing were performed as previously described (Doench et al., 2016; Piccioni et al., 2018). Samples were sequenced on a HiSeq2000 (Illumina). Gene-level analysis was executed using RSA and BAGEL. RSA analysis was conducted as previously described (König et al., 2007). For RSA analysis, DESeq2 was used to generate $\log_2$ fold change from gRNA read counts. Subsequently, z scores were computed in R Studio (version 3.4.2) and RSA scores were generated. For additional significance thresholding, Benjamini Hotchberg correction was performed on RSA values which were plotted against Quantile 3 (Q3) and Quantile 1 (Q1) values.

*Generation and validation of H1 constitutive Cas9 stem cell line:* A targeting vector with AAVS1 homology arms and a Flag-Cas9-2A-Blast-BGHpA expression cassette was generated and co-electroporated with AAVS1 TALENS (System Biosciences) into H1 hESCs using the Neon Transfection System (Thermo Fisher Scientific; Waltham, MA). Two days post-electroporation, Blasticidin (4ug/mL; Thermo Fisher Scientific, R21001) was added and emerging clones were picked and analyzed by immunocytochemistry for FLAG-Cas9 using Mouse anti-FLAG antibody (1:300; Sigma Aldrich, F1804) and by PCR for proper integration into the locus across the junctions (5' junction: *AAVS1-F2* AACTCTGCCCTCTAACGCTG and *CAG-R2* CTATGAACTAATGACCCCGTAATTG; 3' junction: *BGH-F1* GGAAGACAATAGCAGGCATGC and *AAVS1-R4* CCACGTAACCTGAGAAGGGAAT; Non-targeted allele: *AAVS1-F3* CCTGGCCATTGTCACTTTGC and *AAVS1-R4* CCACGTAACCTGAGAAGGGAAT). The H1-36-23 clone was differentiated into neurons using a dual SMAD inhibition protocol and plated into 96-well plates. sgRNAs were delivered by lentiviral vectors that confer puromycin resistance, and the neurons were selected with puromycin (2 μg/mL) for 2 weeks. Neurons were lysed and next-generation sequencing of the gRNA-targeted sites was performed in order to identify and quantify indels generated.

*Validation of primary CRISPR-Cas9 ZIKV survival screen:* SNaPs were induced from H1-36-23 constitutive Cas9 stem cells before lentiviral transduction via spinoculation with individual sgRNAs (pXPR_003 and pXPR_050 vectors) in a 24-well plate format. Cells were then expanded and selected with puromycin (1 μg/mL) for one week. Cas9 gRNA-expressing SNaPs were passaged and plated onto Geltrex at 40,000 cells per well of a 96-well plate (120,000 cells/cm$^2$). Two days later, SNaPs were infected with ZIKV-Ug (MOI = 1) before conducting the infectivity assay at 54 hpi and the cell viability assay at 120 hpi, as previously described. Infectivity and cell viability values were compared to Cas9-SNaPs that were transduced with non-targeting gRNAs.

*Antibody block assay:* SW7338.1 SNaPs were washed once in 1X PBS and then incubated for 1hr at 37°C/5% $CO_2$ in 25 μL of function-inhibiting antibody solution diluted in 1X EBSS: Mouse IgG isotype control (R & D Systems, MAB002) or Mouse anti-integrin aVb5 (EMD Millipore, MAB1961). ZIKV-Ug (MOI = 10) or mock media was then added directly to each well for an additional 1hr at 37°C/5% $CO_2$. Finally, SNaPs were washed once in 1X PBS before being cultured in 100 μL of NPC maintenance media. A 50:50 media exchange was conducted around 24 hpi, and cells were fixed for immunostaining at 54 hpi.

*ZIKV Census-Seq*: Village-44 SNaPs were exposed to mock media or ZIKV-Ug (MOI = 1) for 1 hour at 37°C. After 54 hours, cells were harvested and fixed for 20 minutes at room temperature in BD Cytofix (BD Biosicences, 554714) at a concentration of $1 \times 10^7$ cells/mL. Fixed cells were washed twice with 1X PBS and permeabilized using 1X BD Perm/Wash (BD Biosicences, 554723) at a concentration of $1 \times 10^7$ cells/mL for 10 minutes at room temperature. Cells were then stained with Ms anti-4G2 antibody (1:100) in perm/wash buffer for 1 hr at room temperature in the dark. Stained cells were washed twice with 1X BD Perm/Wash, resuspended in 1X PBS, and kept at 4°C in the dark until analysis. Finally, stained cells were passed through a filter-top 12x75 mm polystyrene tube just before analysis on a BD FACSAria (BD Biosciences; San Jose, CA). Samples were separated based on GFP signal intensity into four bins: ZIKV-Negative (~60% of total cells), ZIKV-Low (~13.3%), ZIKV-Mid (~13.3%), and ZIKV-High (~13.3%).

DNA was unfixed and extracted from each sample using a column-free procedure. First, FAC sorted samples were spun down and resuspended in 300 µL Cell Lysis Solution (Qiagen, 158906) with 2 µL Proteinase K (New England Biolabs, P8107S) and incubated at 56°C overnight. The next day, 1.5 µL of RNase A (Qiagen 158922) was added to each sample prior to a 30 minute incubation at 37°C. Samples were placed on ice for 5 minutes and spun briefly before 200 µL of Protein Precipitation Solution (Qiagen 158910) was added. Samples were vortexed for 20 seconds and then spun down at 13,200 RPM for 10 minutes at 4°C. The supernatant was then transferred to a chilled tube containing 300 µL ice-cold 100% isopropanol with 0.5 µL Glycogen Solution (Qiagen 158930). Samples were again spun down at 13,200 RPM for 10 minutes at 4°C before discarding the supernatant. The DNA pellet was washed in 300 µL of 70% Ethanol and centrifuged at 14,000 RPM for 5 minutes at 4°C. The supernatant was discarded and the DNA pellet was dried at room temperature for 10 minutes so that all of the ethanol was evaporated. Finally, the pellet was re-hydrated at 15-20 µL $dH_2O$ and then incubated for 1 hour at 55°C.

The Census-seq analytical pipeline was executed as previously described (Mitchell et al., 2020). In brief, extracted DNA from FAC-sorted bins were processed for low-coverage DNA sequencing, which was performed using either the TruSeq NanoDNA Library Prep for NeoPrep (Illumina Catalog# NP-101-9001DOC) or Nextera DNA Library Prep (Illumina FC-121-1030) setup (Note: any standard kit that generates sequence libraries from DNA can be compatible with the pipeline). Libraries were then sequenced on an Illumina Nextseq500 instrument using a 75-cycle high output kit. The run was setup as a single 85 bp read and an index read when more than one library was pooled. We regularly pool up to 16 Census-Seq samples in one sequencing run. The output was run through the sequencing alignment protocol using the Picard tools ExtractIlluminaBarcodes and IlluminaBasecallsToSam. The de-multiplexed libraries were then aligned to a human reference genome with BWA. Prior to running the Census-seq algorithm, VCF files were processed to filter variants and add additional site-level information. Variants were first normalized to their appropriate reference sequence using BCFTools. Variants that were monomorphic were dropped, as well as those without a PASS filter, where the site was flagged as problematic during VCF generation. Sites without rsID annotations were updated using information from dbSNP when possible, and otherwise site names were changed to chromosome:position:ref_allele:alt_allele.

Input sequencing and VCF data was then filtered on a per-run basis. Sequence reads were filtered to high quality mappings (MQ>=10) on the autosomes that were not flagged as PCR duplicates. VCF sites were considered if they met all of the following criteria: each site has GQ score of at least 30, is a diploid site, is polymorphic in the subset of donors in the population, and at least 90% of donors have a genotype quality score >=30. In addition, for genotype array-based data where site quality scores may not be available, sites where the reference base is ambiguous [A/T, C/G] were not considered. Only variant sites with ~5% allele frequency were included in analysis. A matrix of donor genotypes and the counts of the reference and alternate allele at each variant were generated. The algorithm initializes with the donor proportions set to equal values (1/number of donors), then runs through an estimation maximization (EM) procedure. The allele frequency of each site is calculated from the genotypes of the donors and their relative proportion in the pool. The initial likelihood of the sequencing data given the starting donor ratios is calculated at each SNP by the likelihood function and the results summed across all sites. To determine how to change the donor ratios to explain the data, an adjustment term is calculated for every donor/site, and the results summed across sites for each donor. This adjustment factor is then scaled by an additional parameter and added to each donor's representation. To determine this scaling value the algorithm employs a univariate optimizer to maximize the donor likelihood. The adjustment is then applied to the data, and the algorithm repeats the adjustment/likelihood optimization loop until convergence.

*Data analysis*: All data were analyzed and plotted using the Prism version 7.03 software (GraphPad; La Jolla, CA) or R 3.5.3.
